# Ras Inhibitor CAPRI Enables Neutrophils to Chemotax Through a Higher-Concentration Range of Gradients

**DOI:** 10.1101/2020.04.23.058131

**Authors:** Xuehua Xu, Xi Wen, Amer Moosa, Smit Bhimani, Tian Jin

## Abstract

Neutrophils sense and migrate through an enormous range of chemoattractant gradients through adaptation. Here, we reveal that, in human neutrophils, Calcium-promoted Ras inactivator (CAPRI) locally controls the GPCR-stimulated Ras adaptation. Human neutrophils lacking CAPRI (*capri^kd^*) exhibit chemoattractant-induced non-adaptive Ras activation; significantly increased phosphorylation of AKT, GSK3α/3β, and cofilin; and excessive actin polymerization. *capri^kd^* cells display defective chemotaxis in response to high-concentration gradients but exhibit improved chemotaxis in low- or subsensitive-concentration gradients of various chemoattractants as a result of their enhanced sensitivity. Taken together, our data reveal that CAPRI controls GPCR activation-mediated Ras adaptation and lowers the sensitivity of human neutrophils so that they are able to chemotax through a higher concentration range of chemoattractant gradients.

**Significance Statement:** Neutrophils provide first-line host defense by migrating through chemoattractant gradients to the sites of inflammation. Inappropriate recruitment and mis-regulated activation of neutrophils contribute to tissue damage and cause autoimmune and inflammatory disease. One fascinating feature of chemotactic neutrophils is their ability to migrate through an enormous concentration range of chemoattractant gradients (10^−9^ ∼ 10^−5^ M) through “adaptation,” in which cells no longer respond to the present stimuli, but remain sensitive to stronger stimuli. The inhibitory mechanism largely remains elusive, although many molecules of the excitatory signaling pathway have been identified. Our study reveals, for the first time, that the inhibitory component, CAPRI, is essential for both the sensitivity and the GPCR-mediated adaptation of human neutrophils.

## Introduction

Neutrophils provide first-line host defense and play pivotal roles in innate and adaptive immunity (1–3). Inappropriate recruitment and dysregulated activation of neutrophils contribute to tissue damage and cause autoimmune and inflammatory diseases (1, 4). Neutrophils sense chemoattractants and migrate to sites of inflammation using G protein-coupled receptors (GPCRs). To accurately navigate through an enormous concentration-range gradient of various chemoattractants (10^−9^ ∼ 10^−5^ M) (Fig. S1), neutrophils employ a mechanism called adaptation, in which they no longer respond to present stimuli but remain sensitive to stronger stimuli. Homogeneous, sustained chemoattractant stimuli trigger transient, adaptive responses in many steps of the GPCR-mediated signaling pathway downstream of heterotrimeric G proteins (5, 6). Adaptation provides a fundamental strategy for eukaryotic cell chemotaxis through large concentration-range gradients of chemoattractants. Abstract models and computational simulations have proposed mechanisms generating the temporal dynamics of adaptation: an increase in receptor occupancy activates two antagonistic signaling processes, namely, a rapid “excitation” that triggers cellular responses and a temporally delayed “inhibition” that terminates the responses and results in adaptation (5, 7–13). Many excitatory components have been identified during last two decades; however, the inhibitor(s) have just begun to be revealed (11, 14–17). It has been recently shown that an elevated Ras activity increases the sensitivity and changes migration behavior (18, 19). However, the molecular connection between the GPCR-mediated adaptation and the cell sensitivity remains missing.

The small GTPase Ras mediates multiple signaling pathways that control directional cell migration in both neutrophils and *Dictyostelium discoideum* (17, 20–24). In *D. discoideum*, Ras is the first signal event that displays GPCR-mediated adaptation (20). Ras signaling is mainly regulated through its activator, guanine nucleotide exchange factor (GEF), and its inactivator, GTPase-activating proteins (GAP) (16, 17, 25). In *D. discoideum*, the roles of DdNF1 and an F-actin-dependent negative feedback mechanism have been previously reported (14, 17). We have previously demonstrated the involvement of locally recruited inhibitors that act on upstream of PI3K in sensing of chemoattractant gradients (11, 26). Recently, we identified a locally recruited RasGAP protein, C2GAP1, that is essential for F-actin-independent Ras adaptation and long-range chemotaxis in *Dictyostelium* (16). Active Ras proteins enrich at the leading edge in both *D. discoideum* cells and neutrophils (17, 27, 28). It has been reported that a RasGEF, RasGRP4, plays a critical role in Ras activation in murine neutrophil chemotaxis (21, 29). However, the components involved in the GPCR-mediated deactivation of Ras and their function in neutrophil chemotaxis are still not known.

In the present study, we show that CAPRI (a <u>Ca</u>lcium-<u>p</u>romoted <u>R</u>as <u>i</u>nactivator) locally controls the GPCR-mediated Ras adaptation in human neutrophils. In response to high-concentration stimuli, cells lacking CAPRI (*capri^kd^*) exhibit non-adaptive Ras activation; significantly increased activation of AKT, GSK3α/3β, and cofilin; excessive actin polymerization; and subsequent defective chemotaxis. Unexpectedly, *capri^kd^* cells display enhanced sensitivity toward chemoattractants and an improved chemotaxis in low- or subsensitive-concentration gradients. Taken together, our findings show that CAPRI Functions as an inhibitory component of Ras signaling, plays a critical role in controlling the concentration range of chemoattractant sensing, and is important for the proper adaptation during chemotaxis.

## Result

### CAPRI regulates GPCR-mediated Ras adaptation in human neutrophils

Chemoattractants induce robust, transient Ras activation in mammalian neutrophils (21, 30). To identify which RasGAP proteins that deactivate Ras in neutrophils, we examined the expression of potential RasGAPs in mouse and human neutrophils (Fig. S2*A*) (31, 32). We found that human and mouse neutrophils highly expressed CAPRI, also called RASA4, consistent with previous reports (33–35). The human neutrophil-like (HL60) cell line provides a useful model to study mammalian neutrophils (22, 36). The differentiated HL60 cell also highly expresses CAPRI (Fig. S2*B*), consistent with a previous report (37), and provides a suitable cell system in which to study CAPRI’s function in human neutrophils. We found that chemoattractant fMLP (N-formyl-L-methionyl-L-leucyl-phenylalanine) stimulation promoted the association between CAPRI and N-Ras/Rap1(Fig 1*A*), suggesting a role of CAPRI in the regulation of chemoattractant-induced Ras and Rap1 signaling. To determine the function of CAPRI, we stably knocked down the expression of *capri* (*capri^kd^*) in HL60 cells using *capri*-specific shRNA lentiviral particles (Fig. 1*B*). We first biochemically determined the dynamics of fMLP-induced Ras activation in both CTL and *capri^kd^* cells using a pull-down assay with a large population of cells (Fig. 1*C*). In resting *capri^kd^* cells, there was a notably higher level of active Ras, indicating CAPRI’s function in regulating basal Ras activity in the cells. In response to 1 μM fMLP stimulation, we detected a transient Ras activation followed by a secondary reactivation in CTL cells as previously reported (14, 16, 30), but a significantly stronger, prolonged Ras activation in *capri^kd^* cells (Fig. 1*D*). We further monitored fMLP-induced Ras activation by visualizing the membrane translocation of a fluorescent active Ras probe, the active Ras binding domain of human Raf1 tagged with RFP (RBD-RFP, red), in single live cells using fluorescence microscopy (Fig. 1*E*). In the present study, we used three plasma membrane (PM) markers, Mem-cerulean, C1AC1A-YFP, and CAAX-mCherry, to track the cell membrane in live cell experiment (Fig. S3). We monitored Ras activation in CTL and *capri^kd^* cells expressing both PM marker (green) and active Ras probe RBD-RFP (red) in response to uniformly applied 1 μM fMLP stimulation. We found that RBD-RFP translocated to and colocalized with PM marker (green) and then returned to the cytoplasm, followed by a second translocation to the protrusion sites of CTL cells (Fig. 1*E*, upper panel, and see Video S1 for a complete, longer time period). In *capri^kd^* cells, the same fMLP stimulation induced persistent membrane translocation and accumulation on the continuously expanding and broadened leading front of the cell (Fig. 1*E*, middle panel, and Video S2). To further examine CAPRI’s function in Ras deactivation, we expressed CAPRI-tagged turboGFP (CAPRI-tGFP), RBD-RFP, and PM marker (Mem-cerulean) in *capri^kd^* cells (*capri^kd^*/^OE^) and monitored fMLP-induced Ras activation in these cells. 1 μM fMLP stimulation triggered a clear membrane translocation of CAPRI-tGFP but much weaker or no membrane translocation of RBD-RFP in *capri^kd^/^OE^* cells (Fig. 1*E*, lower panel, Fig. S4, and Video S3), suggesting that CAPRI translocates to the plasma membrane to inhibit Ras activation (33). Quantitative measurement of RBD-RFP membrane translocation confirmed that fMLP stimulation induced a biphasic Ras activation in CTL cells as previously reported (30), a prolonged Ras activation in *capri^kd^* cells, and a reduced Ras activation in *capri^kd^/^OE^* cells (Fig. 1*F*). Taken together, our results indicate that CAPRI is a RasGAP protein and is required for chemoattractant-mediated Ras adaptation in neutrophils.

**Fig. 1.**
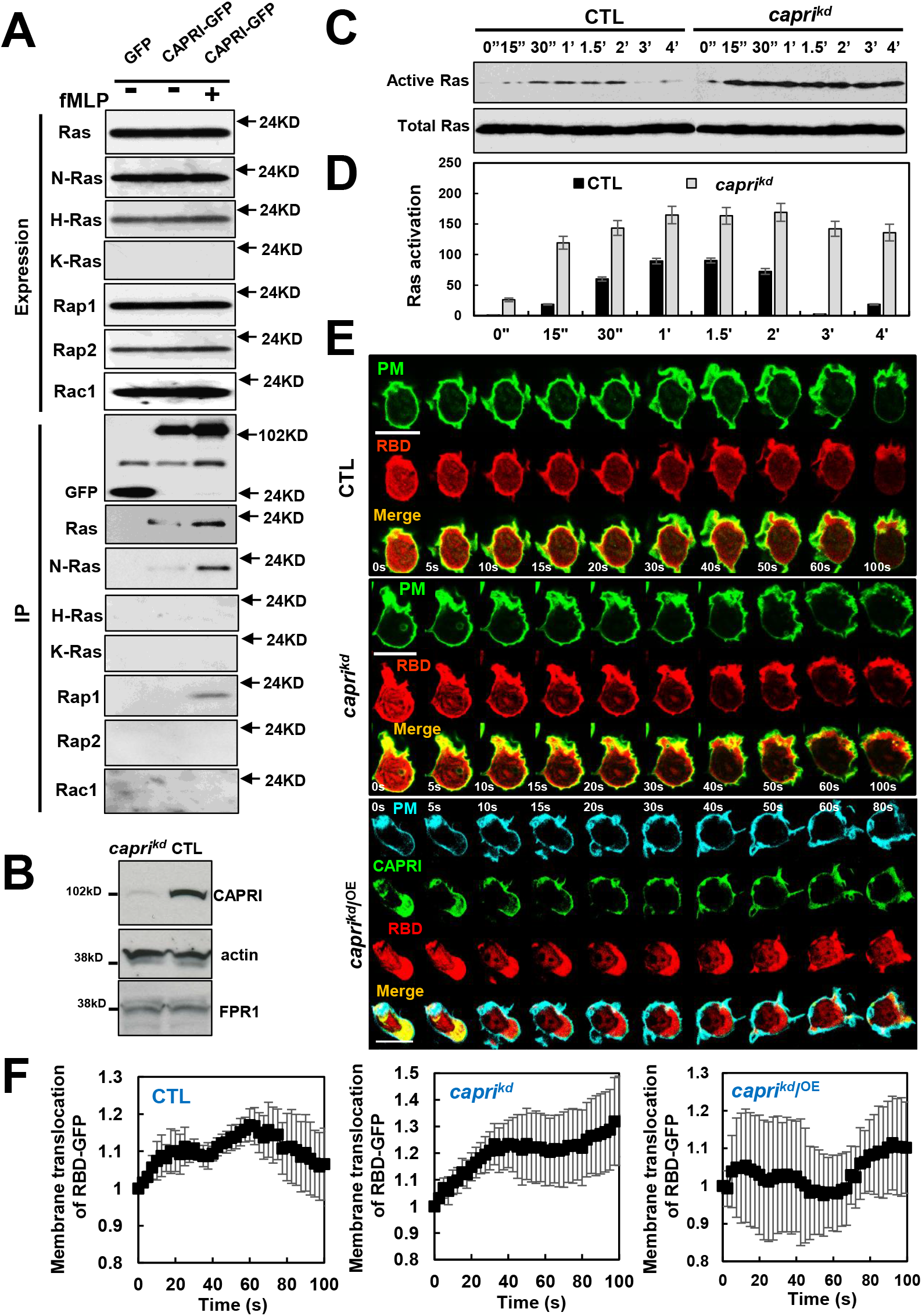
CAPRI interacts with Ras and regulates chemoattractant GPCR-induced Ras adaptation. (A) fMLP-stimulated association between Ras/Rap1/Rac1 and CAPRI detected by co-immunoprecipitation assay. (B) Expression of *capri* in HL60 cells transfected with non-specific (CTL) or *capri* specific (*capri^kd^*) shRNA virus particles. CAPRI was detected by antibodies against human CAPRI. Actin was detected as a loading control; fMLP receptor 1 FPR1 was also detected in both CTL and *capri^kd^* cells. (C) fMLP-induced Ras activation in CTL and *capri^kd^* cells determined by a pull-down assay. Upon stimulation with 10 μM fMLP at time 0”, cells were collected, lysed at the indicated time points, and then centrifuged at 10,000 *g* for 10 min at 4 °C. Agarose beads pre-conjugated with RBD-GST (active Ras binding domain of Raf1 tagged with GST) were incubated with the supernatants of the lysates for 2 hours at 4 °C, then washed with lysis buffer. The protein bound to agarose beads was eluted with 2X SDS loading buffer (SLB). Aliquots of cells at the indicated time points were also mixed with the same volume of SLB for the detection of total Ras protein. The elutes of RBD-GST beads and the aliquots of the cells were analyzed by immunoblotting with anti-pan Ras antibody to detect either active Ras or total Ras protein. (D) Normalized quantitative densitometry of the active Ras from three independent experiments, including the result presented in **C**. The intensity ratio of active Ras and total Ras in WT at time 0 s was normalized to 1. Mean ± SD from three independent experiments is shown. (E) Ras activation in CTL, *capri^kd^*, and *capri^kd^*^/OE^ cells monitored by the membrane translocation of the mRFP-tagged active Ras binding domain of Raf1 (RBD-RFP) in response to 1 μM fMLP stimulation. CTL (top panel and Video S1) and *capri^kd^* (middle panel and Video S2) cells expressed RBD-RFP (red) and a plasma membrane maker (PM, green). The *capri^kd/OE^* cell (bottom panel and Video S3) is a *capri^kd^* cell expressing a PM marker (cyan), CAPRI tagged with turboGFP (CAPRI-tGFP, green), and RBD-RFP (red). fMLP was applied to the cells after 0 s. Scale bar, 10 μm. (F) The quantitative measurement of Ras activation as the membrane translocation of RBD-RFP in E is shown. Mean ± SD is shown; n = 3, 4, and 4 for CTL, *capri^kd^*, and *capri^kd/OE^* cells, respectively.

### Membrane targeting of CAPRI in response to chemoattractant stimulation

To reveal how CAPRI functions during neutrophil chemotaxis, we examined cellular localization of CAPRI-tGFP in HL60 cells. We found that CAPRI-tGFP colocalized with active Ras (RBD-RFP) and actin at the leading edge of chemotaxing HL60 cells in a fMLP gradient (Fig. 2*A*), consistent with previous reports (16, 22, 35). Chemoattractant fMLP and IL-8 induced membrane translocation of CAPRI-tGFP (Fig. 1*E* and Fig. S4-S5). The G protein inhibitor pertussis toxin (PT) blocked the membrane translocation of CAPRI (Fig. S6), indicating that chemoattractant GPCR/G protein-mediated signaling is required for CAPRI membrane targeting. To reveal the molecular mechanism of CAPRI membrane targeting, we investigated the domain requirement for its membrane translocation and interaction with Ras. We expressed tGFP-tagged WT and mutants of ΔC2, ΔPH, and a GAP-inactive mutant R472A, and determined their ability for Ras interaction (Fig. 2*B*-2*C* and Fig. S7). We found that ΔC2 did not interact with Ras and R472A showed a decreased interaction with Ras (Fig. 2*C*), consistent with previous reports (33, 38–41). We next monitored the membrane translocation ability of CAPRI WT and its mutant in response to fMLP stimulation (Fig. 2*D*). Upon fMLP stimulation, WT and R472A clearly translocated to and colocalized with PM marker (Fig. 2*D*, top two panels and Video S4-S5), while ΔPH and ΔC2 showed significantly decreased or little membrane translocation (Fig. 2*D*, lower two panels, and Video S6-S7). Using total internal reflections fluorescent (TIRF) microscopy, we confirmed the membrane-translocation behavior of CAPRI WT and its mutants (Fig. S8 and Fig. 2*E*) (42). Our results indicate that both the C2 and PH domains are crucial for chemoattractant-induced membrane targeting.

**Fig. 2.**
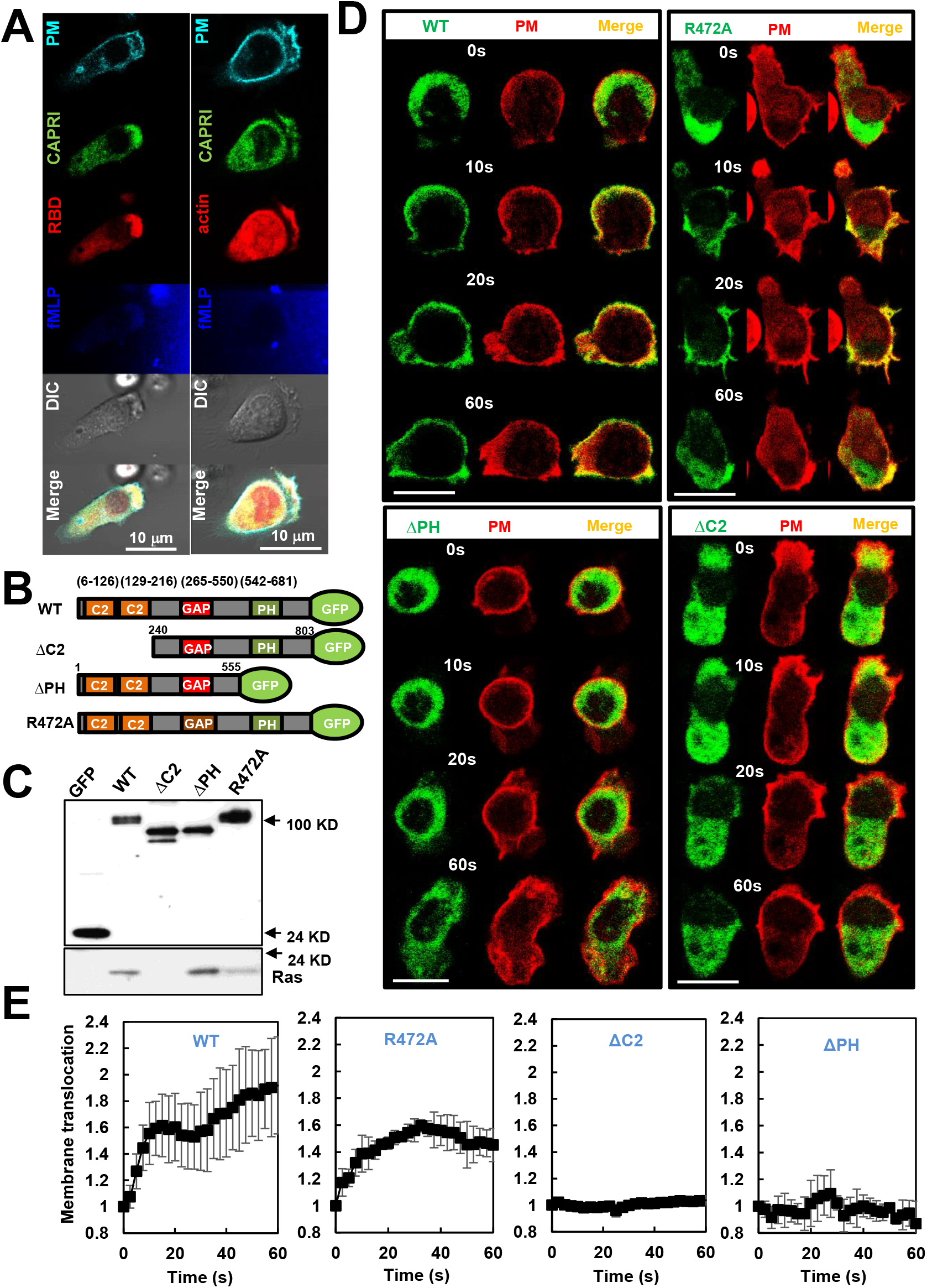
Chemoattractant-induced membrane targeting of CAPRI. (A) Colocalization of active Ras and Rac1 with CAPRI in the leading edge of a chemotaxing cell. HL60 cells expressed a PM marker (cyan), both tGFP-tagged CAPRI (green), and RBD-RFP (red) or actin-mCherry (red) were exposed to a 100 nM fMLP gradient (dark blue). To visualize the fMLP gradient, 100 nM fMLP was mixed with Alexa 633 (dark blue). Scale bar = 10 μm. (B) The scheme shows the domain composition of tGFP-tagged wild-type (WT) or mutants of CAPRI. (C) Domain requirement for the interaction of CAPRI and Ras determined by co-immunoprecipitation. HL60 cells expressing either tGFP alone or tGFP-tagged WT or mutants of CAPRI were lysed with immunoprecipitation buffer with 1 mM GTPγS and went through the coimmunoprecipitation process and western blotting using anti-GFP or Ras antibodies. (D) Montage shows the membrane translocation of CAPRI and its mutants upon uniform application of 1 μM fMLP. Cells expressing tGFP-tagged WT or mutants of CAPRI (green) and PM marker (red) were imaged in time-lapse and 100 nM fMLP (red) was applied to the cells after 0 s. Scale bar = 10 μm. Videos S4-S7 are CTL cells expressing PM marker (red) and tGFP-tagged WT, R472A, ΔPH or ΔC2 of CAPRI (green), respectively. (E) The quantitative measurement of the membrane translocation of CAPRI or its mutants is shown. Mean ± SD is shown, n = 5, 4, 4, and 4 for WT, ΔC2, ΔPH, or R472A of CAPRI, respectively. The quantitative measurement of intensity changes was described in the Materials and Methods section.

### Chemoattractant induces hyperactivation of Ras effectors in *capri^kd^* cells

PI3Kγ is a direct effector of Ras, which synthesizes lipid phosphatidylinositol (3,4,5)-trisphosphate (PtdIns(3,4,5)P_3_, PIP_3_) and activates the PIP_3_-binding protein AKT in neutrophils (21, 29, 43). To examine the consequence of non-adaptive Ras activation, we monitored fMLP-induced PIP_3_ production using a biosensor PH-GFP (PH-domain of human AKT tagged with GFP, green) in both CTL and *capri^kd^* cells (Fig. 3*A*) (44). We found that 1 μM fMLP trigged a robust translocation of PH-GFP to the entire plasma membrane followed by a partial return to the cytosol and a gradual accumulation in the protrusion site of CTL cell (Fig. 3*A*, upper panel, and Video S8). The same stimulation induced a stronger and persistent accumulation of PH-GFP all around the periphery of *capri^kd^* cells (Fig. 3*A*, lower panel, and Video S9), indicating a hyperactivation of PI3K in cells lacking CAPRI. We further determined the activation profile of PI_3_Kγ downstream effectors in CTL and *capri^kd^* cells. AKT and GSK are well-known effectors of Ras/PI3Kγ signaling and are critical for the reorganization of actin cytoskeleton in neutrophil chemotaxis (22, 45). Therefore, we examined fMLP-induced PI3Kγ activation by measuring the phosphorylation of AKT on residues T308 and T473 in both CTL and *capri^kd^* cells. We found that 1 μM fMLP triggered a transient phosphorylation on T308 and T473 of AKT in CTL cells, while it induced a persistent and significantly increased phosphorylation on both residues of AKT in *capri^kd^* cells (Fig. 3*B*-3*C*), consistent with previous reports (46–48). Cofilin, an F-actin depolymerization factor (ADF), is essential for depolymerization of F-actin in dynamic reorganization of the actin cytoskeleton during cell migration (49). Its activity is regulated mainly through a phosphorylation event: phosphorylation on Ser-3 inhibits its actin binding, severing, and depolymerizing activities; and dephosphorylation on Ser-3 by slingshot proteins (SSHs) reactivates it. Slingshot 2 is a direct substrate of GSK3 and chemoattractants induce phosphorylation and inhibition of GSK3α/3β partially through PI3Kγ-AKT pathways in neutrophils (45). As previously reported (45), GSK3α/3β is constitutively active and dephosphorylated in resting cells, and fMLP induced a transient phosphorylation and deactivation of GSK3α/3β and led to a transient dephosphorylation of cofilin (Fig. 3*D*-3*E*). We found that 1 μM fMLP stimulation triggered significantly stronger, prolonged phosphorylation of GSK3α/3β and persistent dephosphorylation of cofilin in *capri^kd^* cells. Taken together, these results indicate that CAPRI is essential for the proper activation/deactivation dynamics of the PI3Kγ/AKT/GSK3/cofilin signaling pathway in response to chemoattractant stimulations.

**Fig. 3.**
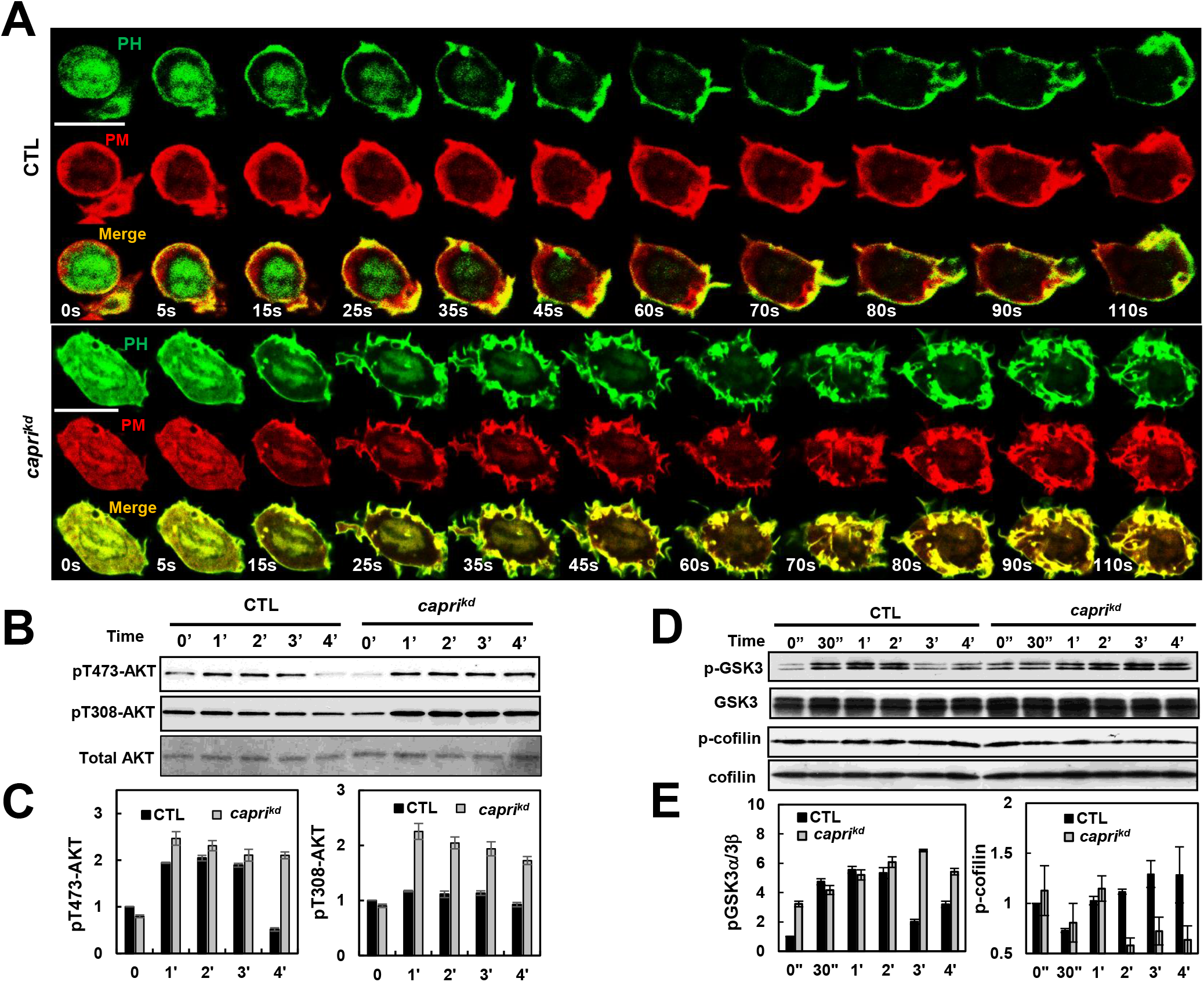
Enhanced chemoattractant-induced activation of PI3K signaling in *capri^kd^* cells. (A) Montage shows the membrane translocation of PIP_3_ biosensor PH-GFP in CTL and *capri^kd^* cells in response to 1 μM fMLP stimulation. Cells expressing PH-GFP (GFP-tagged PH domain of human AKT, green) and PM marker (red) were stimulated with 1 μM fMLP at time 0 s. Scale bar = 10 μm. Videos S8 and S9 are CTL and *capri^kd^* cells, respectively. (B) fMLP-induced phosphorylation of AKT in CTL and *capri^kd^* cells. A final concentration of 10 μM fMLP stimulation was added to the cells at time 0 s. Aliquots of cells were sampled at the indicated time points and subjected to western blot detection of the phosphorylated and total proteins of interest in **A** and **C**. (C) Normalized quantitative densitometry of phosphorylated AKT by total AKT protein in **B**. Mean ± SD of three independent experiments is shown. The intensity ratio of the phosphorylated versus total AKT in CTL cells at time 0 s is normalized to 1. (D) fMLP-induced phosphorylation of GSK3α/3β and cofilin in CTL and *capri^kd^* cells. (E) Normalized quantitative densitometry of the phosphorylated GSK3α/3β, and cofilin in **D**. The intensity ratio of the phosphorylated versus total GSK3α/β and cofilin in CTL cells at time 0 s is normalized to 1. Mean ± SD of three independent experiments is shown.

### fMLP stimulation induces an increased activation of Rap1 and its effector in *capri^kd^* cells

Rap1 is a close relative of Ras initially described as a competitor of Ras by directly interacting with Ras effectors (50, 51). Later studies indicate that Rap1 also functions in independent signaling pathways to control diverse processes, such as cell adhesion, cell-cell junction formation, and cell polarity (52, 53). It has been shown that CAPRI also interacts with Rap1 in CHO cells (54). We found that fMLP stimulation promoted the interaction between CAPRI and Rap1 in neutrophils (Fig. 1*A*) and that CAPRI was recruited to adhesion sites and colocalized with the PM marker during cell migration (Fig. S9). To understand CAPRI’s function in Rap1 activation in human neutrophils, we determined fMLP-induced Rap1 activation in both CTL and *capri^kd^* cells by a pull-down assay using the Rap-GTP binding domain of human RalGDS (RBD_RalGDS_) (Fig. 4*A*). There was a higher level of active Rap1 in resting *capri^kd^* cells. fMLP stimulation (1 μM) induced a transient Rap1 activation in CTL cells, while it triggered a significantly increased and prolonged Rap1 activation in *capri^kd^* cells (Fig. 4*B*). We further monitored Rap1 activation using an active Rap1 probe, GFP-tagged RBD_RalGDS_ (RBD_RalGDS_-GFP) (52), using live cell fluorescence microscopy (Fig. 4*C*). We found that RBD_RalGDS_-GFP colocalized with PM marker at the adhesion/protrusion sites of the resting cells (Fig. 4*C*), suggesting a potential function of Rap1 in the adhesion of neutrophils. Upon 1 μM fMLP stimulation, RBD_RalGDS_-GFP (green) transiently translocated to and colocalized with PM marker (red), then returned to the cytoplasm, and then translocated to and colocalized with the PM marker again at the protruding sites of CTL cells (Fig. 4*C*, upper panel, and Video S10). In resting *capri^kd^* cells, there was notably more membrane localization of RBD_RalGDS_-GFP, which is consistent with the notion that there is a higher basal Rap1 activity in *capri^kd^* cells (Fig. 4*C*, lower panel). We further found that 1 μM fMLP triggered prolonged translocation of RBD_RalGDS_-GFP (green) to and colocalization with the PM marker (red) during a continuous expansion of *capri^kd^* cells (Video S11), suggesting that the fMLP induces a hyperactivation of Rap1 in cells lacking CAPRI. To understand the effects of Rap1 hyperactivation, we examined Rap1 effector Erk42/44 phosphorylation (Fig. 4*D*). We detected higher basal Erk42/44 phosphorylation in the non-stimulated *capri^kd^* cells. In contrast to a rapid maximum phosphorylation of Erk 42/44 at 30 s in CTL cells, same stimulation induced an increasing dynamics of Erk42/44 phosphorylation in *capri^kd^* cells (Fig. 4*E*), consistent with previous reports (33, 35, 54). Our results suggest that CAPRI deactivates Rap1 activation to facilitate an appropriate Rap1 activation and its effector for chemotaxis in neutrophils.

**Fig. 4.**
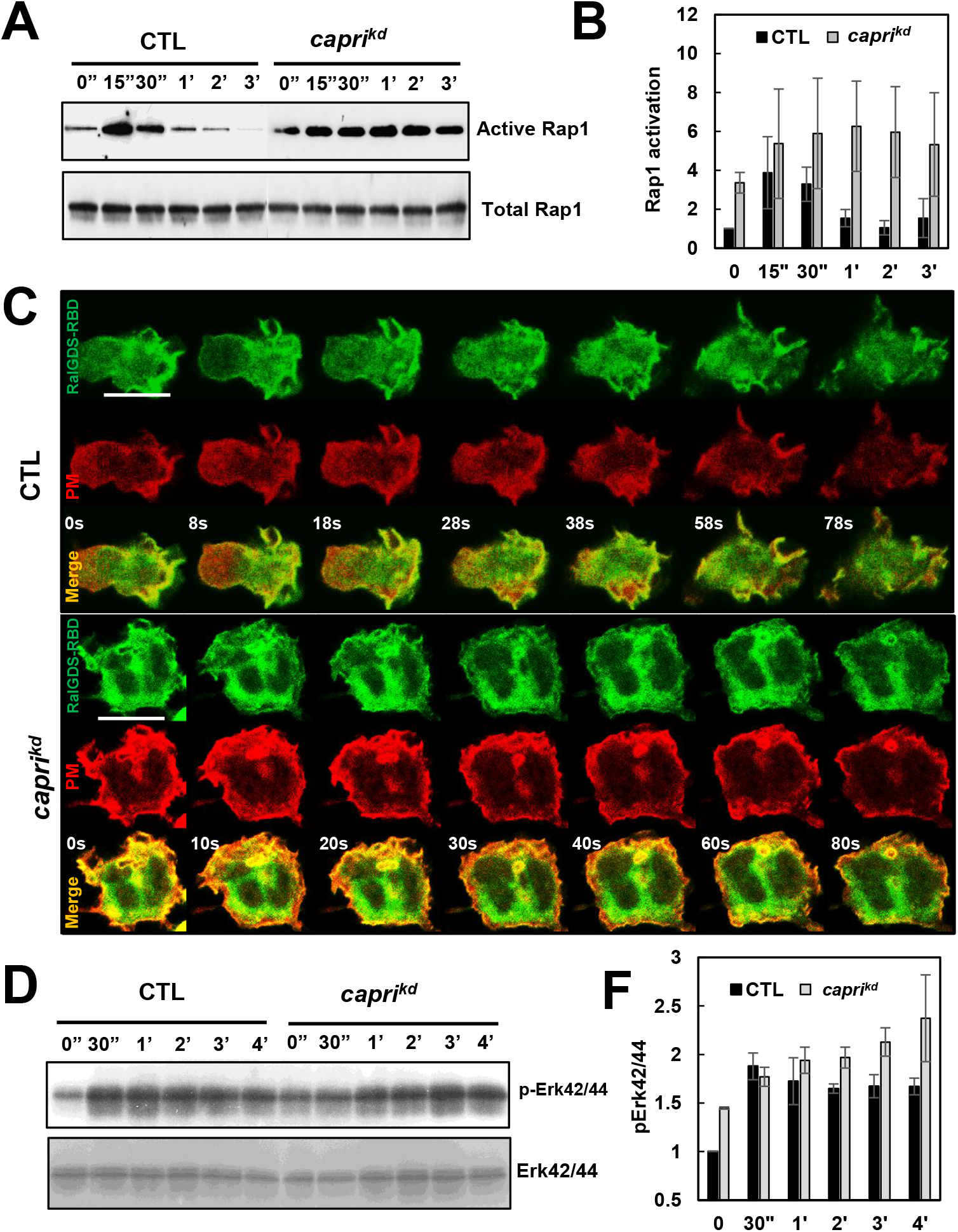
Increased activation of Rap1 and its effector induced by fMLP stimulation in *capri^kd^* cells. (A) fMLP-induced Rap1 activation in CTL and *capri^kd^* cells determined by a pull-down assay using GST-RBD_RalGDS_ agarose beads. (B) Normalized quantitative densitometry of the active Rap1 from three independent experiments, including the result presented in A. The intensity ratio of active Rap1 and total Rap1 in WT at time 0 s was normalized to 1. Mean ± SD from three independent experiments is shown. (C) fMLP-induced Rap1 activation monitored by the membrane translocation of the active Rap1 probe GFP-RBD_RalGSD_. Cells expressing RBD_RalGSD_-GFP (green) and PM marker (red) were stimulated with 1 μM fMLP at 0 s. Scale bar = 10 μm. Videos S11-S12 are CTL and *capri^kd^* cells, respectively. (D) fMLP-induced phosphorylation of Erk42/44 in CTL and *capri^kd^* cells. (E) Normalized quantitative densitometry of the phosphorylated Erk42/44 in E. The intensity ratio of the phosphorylated versus total Erk42/44 in CTL cells at time 0 s is normalized to 1. Mean ± SD of three independent experiments is shown.

### Excessive polymerization of actin impairs the polarization and migration of *capri^kd^* cells

The chemoattractant GPCR/G-protein signaling regulates spatiotemporal activities of Ras and Rap1 that mediate multiple signaling pathways to control the dynamics of actin cytoskeleton that drives cell migration. To evaluate the role of CAPRI in chemoattractant GPCR-mediated actin assembly in neutrophils, we determined fMLP-mediated polymerization of actin in CTL and *capri^kd^* cells using a centrifugation assay of actin filaments (F-actin) (Fig. 5*A*). In CTL cells, 10 μM fMLP stimulation induced a transient polymerization of actin. In *capri^kd^* cells, the same stimulation also triggered the initial, transient actin polymerization followed by a much stronger and persistent actin polymerization (Fig. 5*A*-5*B*). To understand the temporospatial dynamics of actin polymerization, we next monitored actin polymerization using the membrane translocation of an actin filament probe, F-tractin-GFP, in live cells by fluorescence microscopy (55). We found that, in response to uniformly applied 1 μM fMLP, F-tractin-GFP (green) translocated to and colocalized with the PM marker (red) along the entire periphery of CTL cells, then withdrew, and accumulated at the protruding sites of CTL cells (Fig. 5*C*, upper panel and Video S12), indicating an initial overall actin polymerization on the entire periphery followed by a localized polymerization on the protrusion sites. In *capri^kd^* cells, the same stimulation trigged a stronger and prolonged membrane translocation of F-tractin-GFP, followed by a slight withdrawal, and then continuous accumulation on multiple expending/ruffling sites of cells (Fig. 5*C*, lower panel and Video S13). To understand the effect of excessive actin polymerization on chemotaxis, we visualized the distribution of F-actin in cells migrating toward a 1 μM fMLP gradient using another F-actin probe (SiR-actin, a live-cell staining probe) (Fig. 5*D*). As expected, CTL cells displayed a clearly polarized actin polymerization: a protruding leading front (pseudopod) and a contracting trailing edge (uropod), and they chemotaxed through the gradient and accumulated at the source of the fMLP gradient (Fig. 5*D*, left panel and Video S14). However, the *capri^kd^* cells, especially those close to the source of the 1 μM fMLP gradient, displayed an overall excessive F-actin distribution, poor polarization, and significantly slow migration during chemotaxis (Fig. 5*D*, right panel and Video S15). Notably, some of the *capri^kd^* cells located relatively far away from the source of fMLP showed polarized morphology and migrated effectively toward the fMLP source, but gradually lost their polarity and stopped moving when they were close to the source of the fMLP (Video S15). The above results together indicate that chemoattractant stimuli at high concentrations induce excessive polymerization of actin that impairs the polarization and migration of *capri^kd^* cells.

**Fig. 5.**
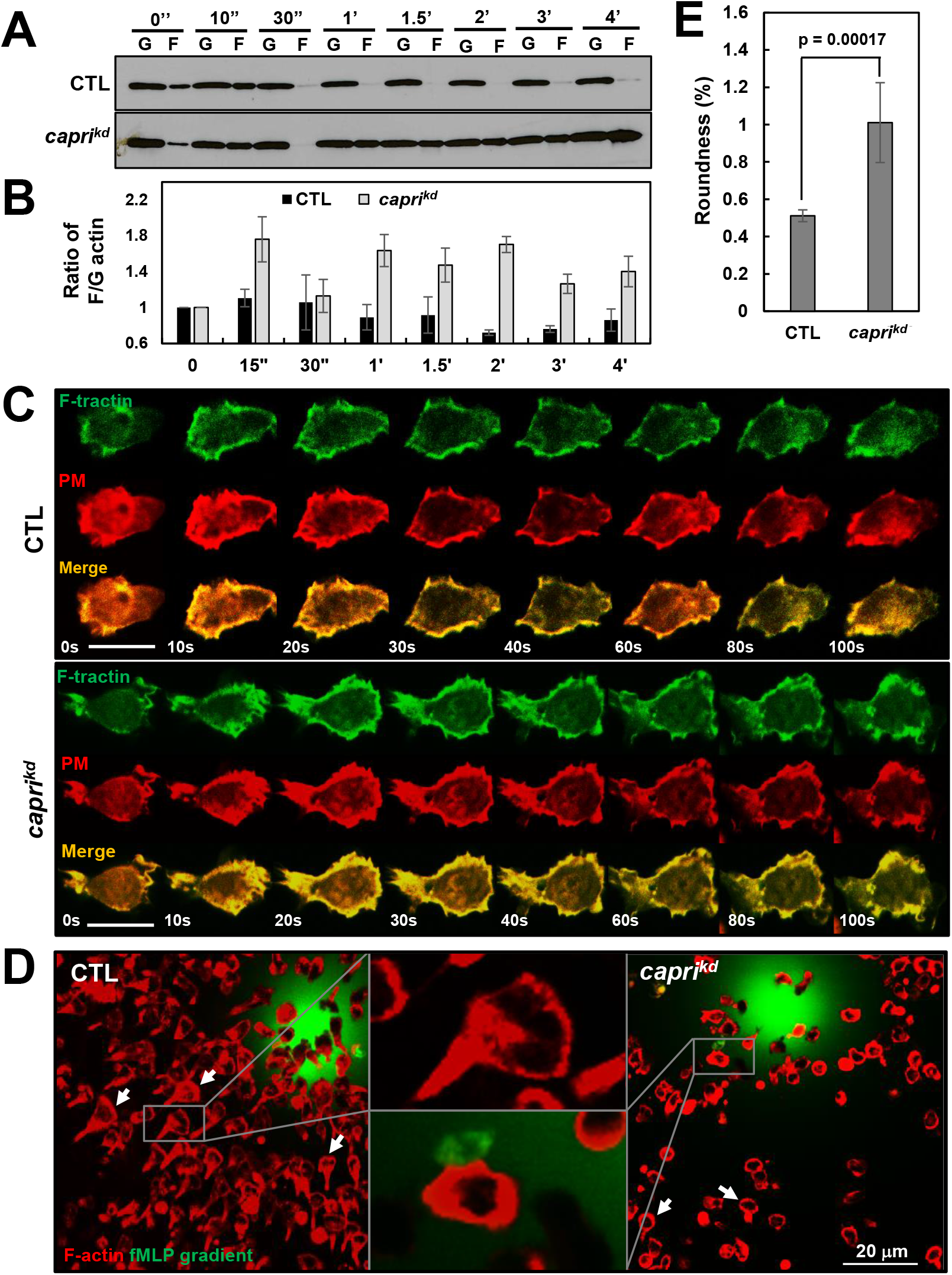
Elevated polymerization of actin in response to a high concentration of fMLP stimulation impairs the polarity and migration of *capri^kd^* cells. (A) The amount of globular (G) and filamentous (F) actin in CTL and *capri^kd^* cells was determined by a centrifugation assay of F-actin. Cells were stimulated with 10 μM fMLP at time 0 s and aliquots of cells at the indicated time points were analyzed. (B) Normalized quantitative densitometry of the F/G-actin ratio in the CTL and of *capri^kd^* cells in (A). Mean ± SD from three independent experiments is shown. The F/G ratio of CTL cells at 0 s was normalized to 1. (C) fMLP-induced actin polymerization by monitoring F-actin probe GFP-F-tractin using live cell confocal microscopy in CTL and *capri^kd^* cells. Cells expressing tractin-GFP (green) and PM marker (red) were stimulated with 1 μM fMLP at time 0 s. Scale bar = 10 μm. Videos S12-S13 are CTL and *capri^kd^* cells, respectively. (D) F-actin distribution in chemotaxing CTL and *capri^kd^* cells in a 1 μM fMLP gradient. The differentiated cells were stained with the actin filament probe, SiR-actin (red). Cells were exposed to a 1 μM fMLP gradient and allowed to chemotax for 5 min. Arrows point to the cells with a polarized, chemotaxing morphology. To visualize the gradient, fMLP was mixed with Alexa 488 (green, 1 μg/ml). Scale bar = 20 μm. Videos S14-S15 are CTL and *capri^kd^* cells, respectively. (E) The graph shows the roundness of CTL and *capri^kd^* cells shown in **D**. The roundness was measured as the ratio of the width vs the length of the cell. That is, the roundness for a circle is 1 and for a line is 0. Student’s *t*-test was used to calculate the *p* value.

### *capri^kd^* cells display improved chemotaxis in low- or subsensitive-concentration gradients of chemoattractants, but defective chemotaxis in high-concentration gradients

To further determine the function of CAPRI in chemotaxis of neutrophils, we monitored chemotaxis behavior of CTL and *capri^kd^* cells in gradients of fMLP at different concentrations using an *EZ-TAXIScan* analysis (56). In a 1 μM fMLP gradient, *capri^kd^* cells, in a clear contrast to CTL cells, displayed a significant decrease in migrating speed (CTL: 20.86 ± 3.12 μm/min; *caprikd:* 12.16 ± 3.73 μm/min), directionality (CTL: 0.86 ± 0.09; *caprikd:* 0.73 ± 0.14 μm/min), total path length (CTL: 140.56 ± 27. 7 μm; *caprikd:* 86.74 ± 13.75 μm), and polarity measured as roundness of a cell (%) (CTL: 76.2 ± 6.81; *caprikd:* 84.82 ± 5.76) (Fig. 6*A*-6*B*). To understand chemotaxis behavior over a large concentration range, we further monitored the chemotaxis behavior of both CTL and *capri^kd^* cells in the fMLP gradients at lower concentrations (Fig. 6*C*). We found that severe defects in chemotaxis in *capri^kd^* cells were clearly observed when they experienced fMLP gradients at high concentration (>100 nM). When exposed to a gradient generated from a lower concentration (10 nM fMLP source, both CTL and *capri^kd^* cells displayed similar directionality, although *capri^kd^* cells still displayed decreased speed and total path length (Fig. 6*D*). In response to a gradient generated from a 1 nM fMLP source, *capri^kd^* cells displayed significantly better directionality, but similar migration speed, in comparison with CTL cells. In a 0.1 nM fMLP gradient, most CTL cells displayed random migration, while the majority of *capri^kd^* cells displayed improved directionality and migration speed. Without a gradient, *capri^kd^* cells displayed a bigger random walk compared to CTL cells. That is, *capri^kd^* cells display defective chemotaxis in high-concentration gradients but improved chemotaxis in low- or subsensitive-concentration gradients of chemoattractants. We also observed this concentration-dependent change of chemotaxis capability of *capri^kd^* cells in response to gradients of IL-8, another chemoattractant for neutrophils (Fig. S10). The above result indicates that CTL cells chemotaxed efficiently through gradients of various chemoattractants ranging from 10^−9^ to 10^−6^ M, while *capri^kd^* cells did so from 10^−10^ to 10^−7^ M.

**Fig. 6.**
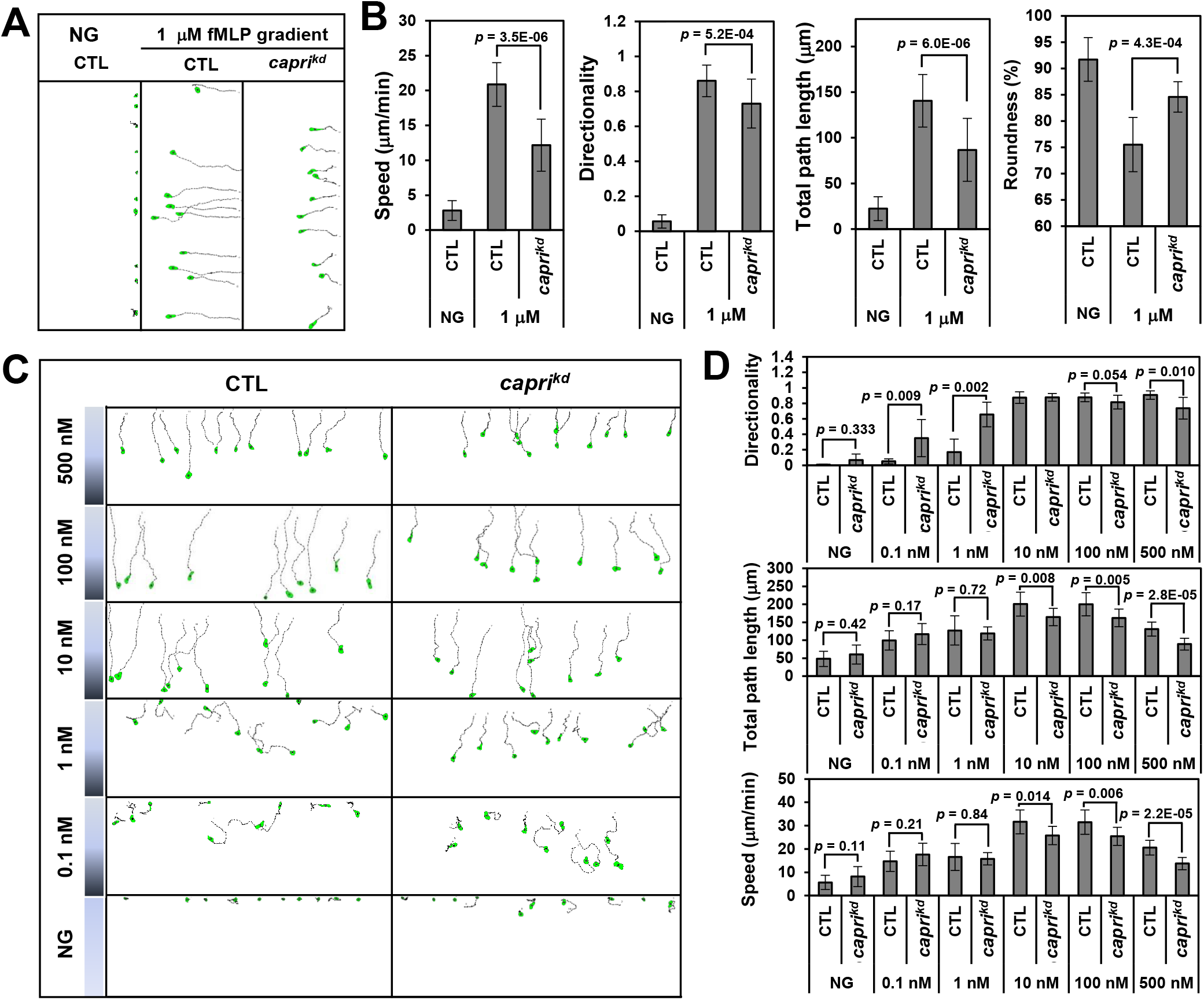
The concentration range of fMLP gradients in which neutrophils chemotax efficiently is upshifted in *capri^kd^* cells. (A) Montages showing the travel path of chemotaxing CTL or *capri^kd^* cells in a gradient generated from a 1 μM fMLP source. Cell movement was analyzed by DIAS software. (B) Chemotaxis behaviors measured from A are described as four parameters: directionality, which is “upward” directionality, where 0 represents random movement and 1 represents straight movement toward the gradient; speed, defined as the distance that the centroid of the cell moves as a function of time; total path length, the total distance the cell has traveled; and roundness (%) for polarization, which is calculated as the ratio of the width to the length of the cell. Thus, a circle (no polarization) is 1 and a line (perfect polarization) is 0. Thirteen cells from each group were measured and the mean ± SD was shown. Student’s *t*-test was used to calculate the *p* value. (C) Montages show the travel path of chemotaxing CTL or *capri^kd^* cells in response to a wide range of fMLP gradients. (D) Chemotaxis behaviors measured from **C** are described by three parameters: directionality, speed, and total path length as described in **B**. Thirteen cells from each group were measured. Student’s *t*-test was used to calculate the *p* value.

### An increased sensitivity of *capri^kd^* cells toward chemoattractants

Using TIRF imaging, we noticed that CAPRI-tGFP localized on the membrane of resting cells, suggesting its potential role in inhibiting basal Ras activity (Fig. 7*A*). As expected, fMLP stimulation induced a strong, quick membrane translocation of CAPRI followed by a slow, gradual withdrawal with a notable fraction of CAPRI remaining on the plasma membrane for quite a long period of time. We also observed a higher basal Ras activity in *capri^kd^* cells (Fig. 1*C*), indicating a potential role of CAPRI in maintaining a low basal Ras activity in the resting cells. The basal activity of Ras is part of a positive feedback mechanism that promotes actin dynamics and, more importantly, chemotaxis in a shallow chemoattractant gradient in *D. discoideum* (24). We speculated that the higher basal Ras activity in *capri^kd^* cells might contribute to their enhanced random walk without a gradient or to their improved chemotaxis in a low- or subsensitive-concentration gradient (Fig. 6). To determine whether *capri^kd^* cells have become more sensitive to the stimulus, we determined Ras and Rap1 activation in response to fMLP stimulation at different concentrations using a pull-down assay (Fig. 7*B*). Comparing to the CTL cells, we detected a stronger Ras/Rap1 activation in *capri^kd^* cells in response to all three different concentrations (Fig. 7*C*). In response to 0.1 nM fMLP stimulation, CTL cells did not show a clear Ras or Rap1 activation but displayed a weak oscillation, while *capri^kd^* cells showed a clear transient Ras activation. In response to 10 nM and 1 μM fMLP stimulation, *capri^kd^* cells showed a stronger, longer activation of both Ras and Rap1 than CTL cells did.

**Fig. 7.**
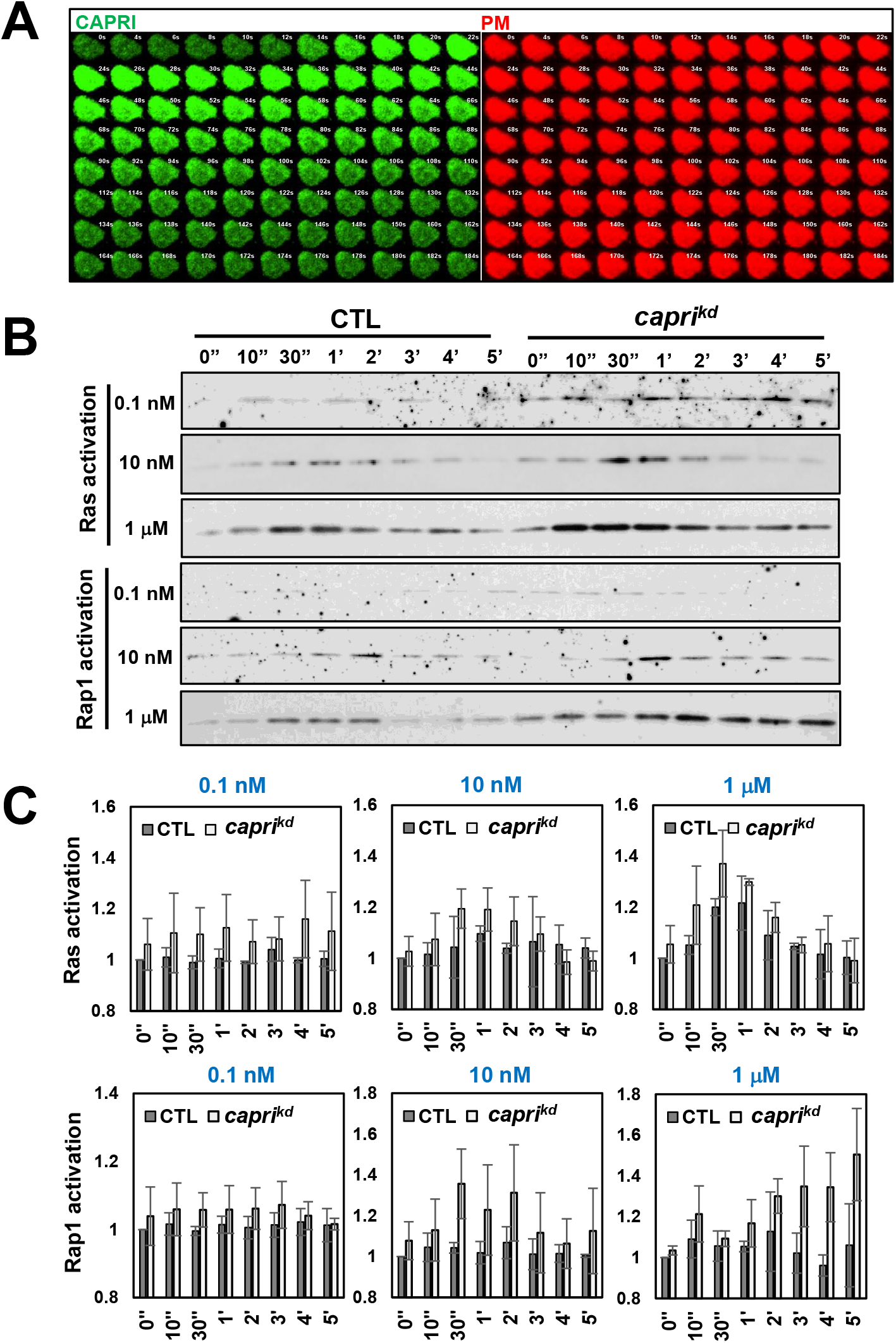
Activity of Ras and Rap1 is increased in *capri^kd^* cells. (A) CAPRI localizes at the plasma membrane before and after fMLP stimulation. (B) Ras and Rap1 activation in CTL and *capri^kd^* cells in response to 0.1 nM, 10 nM, or 1 μM fMLP stimulation determined by a pull-down assay. (C) Normalized quantitative densitometry of the active Ras and Rap1 from three independent experiments, including the result presented in **B**. The intensity ratio of active Ras and Rap1 in WT at time 0 s was normalized to 1. Mean ± SD from three independent experiments is shown.

We further determined the responsiveness of both CTL and *capri^kd^* cells to either 0.1 nM or 10 nM fMLP stimulation by monitoring actin polymerization through the membrane translocation of F-tractin-GFP (Fig. 8). In response to 0.1 nM fMLP stimulation, most CTL cells (∼ 90 %) did not show clear membrane translocation of F-tractin-GFP to the plasma membrane, while they showed a continuous translocation of F-tractin-GFP to the protrusion sites (Fig. 8*A*, top panel, and Video S16). In contrast, more than 80 % of the *capri^kd^* cells showed clear membrane translocation of F-tractin upon 0.1 nM fMLP stimulation (Video S17 and Fig. 8*B*), indicating a higher sensitivity of *capri^kd^* cells. Not surprisingly, 10 nM fMLP induced robust membrane translocation of F-tractin in both CTL and *capri^kd^* cells with different dynamics: CTL cells showed a clear transient translocation of F-tractin-GFP to plasma membrane (∼ 5-40 s), followed by a withdrawal of most F-tractin from the plasma membrane (∼ 50-60 s) and then a second translocation to the protrusion sites (> 60 s) (Video S18); while *capri^kd^* cells showed a stronger and longer membrane translocation of F-tractin-GFP to plasma membrane (∼ 5-90 s) followed by a gradual withdrawal (∼ 90-130 s) (Video S19), indicating a stronger response and a longer period of time required to adapt the stimulation. Consistent with the results above, a prolonged, non-adaptive actin polymerization was observed in *capri^kd^* cells in response to stimuli at higher concentrations (Fig. 5 and Fig. 8*C*). These results together demonstrate that *capri^kd^* cells, lacking Ras inhibitor, are sensitive to subsensitive-concentration chemoattractant stimuli for CTL cells, while they fail to chemotax through a high-concentration gradient as CTL cells do.

**Fig. 8.**
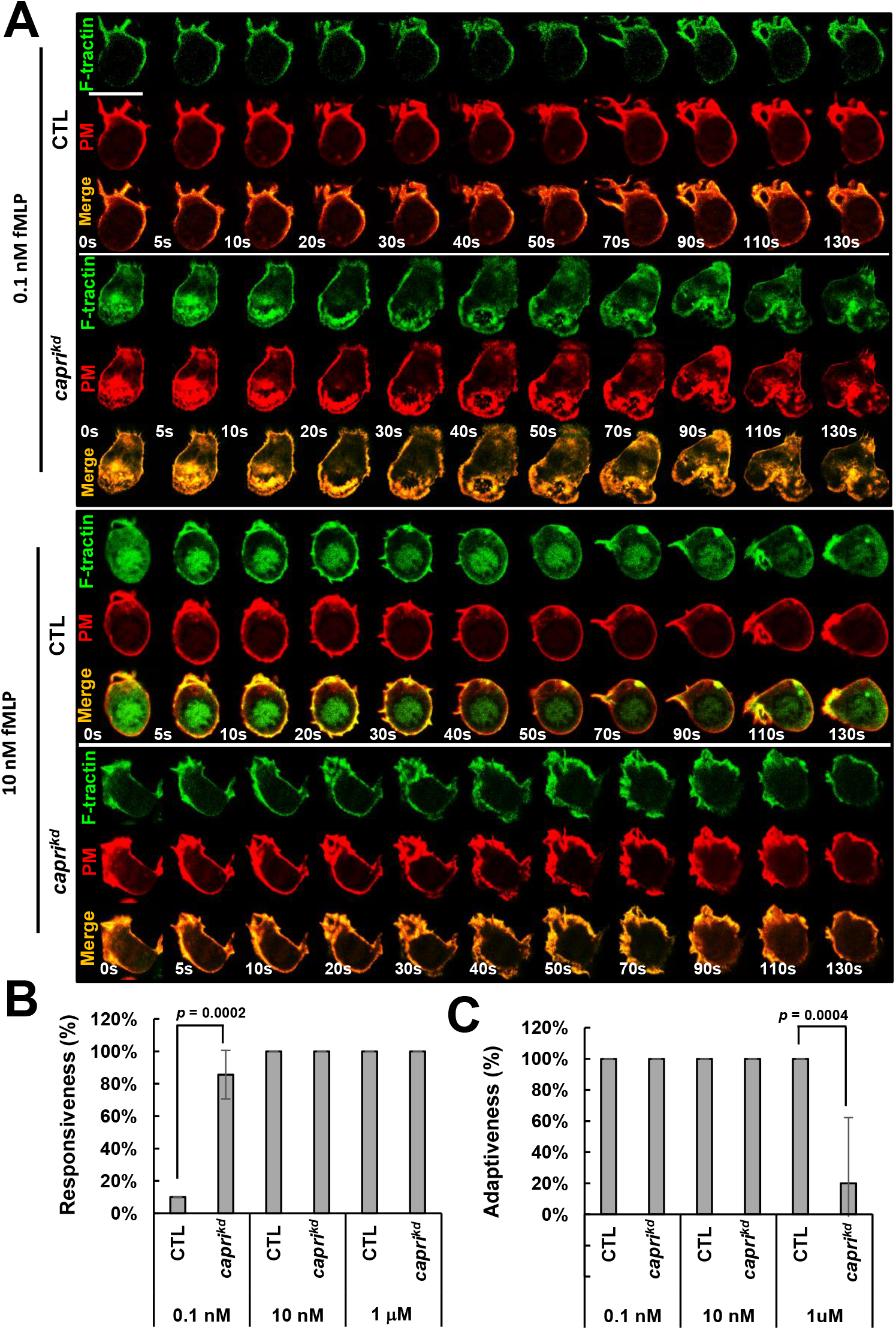
*capri^kd^* cells are more sensitive to the stimuli at low concentration and fail to adapt to the stimuli at high concentrations. (A) Montage shows cell response as actin polymerization monitored through the membrane translocation of F-actin probe, Ftractin-GFP, in both CTL or *capri^kd^* HL60 cells using fluorescent microscopy. Cells expressing GFP-F-tractin (green) and PM marker (red) were stimulated with either 0.1 nM or 10 nM fMLP stimulation. Scale bar = 10 μm. Videos S16 and S18 are CTL cells and Videos S17 and S19 are *capri^kd^* cells. Videos S16-S17 and Videos S18-19 were cells stimulated with either 0.1 nM or 10 nM fMLP, respectively. (B) The percentage of cells that respond to the indicated concentration of fMLP stimuli in CTL and *capri^kd^* cells is shown. (C) Percentage of cells with adaptation in response to fMLP stimulation. **B** and **C** were measured with the same sets of data. The numbers of independent experiments for CTL and *capri^kd^* cells for 0.1 nM, 10 nM, and 1 μM fMLP stimuli are 11 and 7, 8 and 10, and 5 and 9, respectively. Student’s *t*-test was used to calculate the *p* value.

## Discussion

Here, we show that CAPRI is a locally membrane-recruited negative regulator of Ras signaling and enables neutrophils to chemotax through a higher concentration range of chemoattractant gradients by lowering their sensitivity and through GPCR-mediated adaptation.

### Chemoattractant-induced deactivation of Ras is regulated through the local membrane recruitment of CAPRI

The membrane recruitment of CAPRI and its function in Ras deactivation have been previously studied (33, 39). CAPRI, CAlcium-<u>P</u>romoted <u>R</u>as <u>I</u>nactivator, was first characterized by its calcium-dependency on deactivation of Ras (33). In this study, we found that the ΔC2 mutant of CAPRI did not translocate in response to chemoattractant stimulation, indicating an essential role of C2/calcium binding in the membrane targeting of CAPRI (Fig. 2*E*). In addition to the C2 domain, we found that the ΔPH mutant showed significantly decreased membrane translocation in response to chemoattractant stimulation (Fig. 2*D*-2*E*). This result is consistent with the previous reports that the PH domain of the GAP1 family is responsible for, or plays an important role, in their membrane translocation (34, 39, 57, 58). We also found that deactivation of GAP activity of the GAP domain (R472A mutant) had no effect on CAPRI’s membrane targeting (Fig. 2*D*), although it might affect its interaction with Ras (Fig. 2*E*), consistent with the notion that the arginine residue of the GAP domain not only is responsible for its GAP activity, but also stabilizes its interaction with Ras (41). We also found that *capri^kd/OE^* cells, which displayed little or no Ras activation, still exhibited a strong CAPRI membrane translocation, indicating that active status of Ras is not required for CAPRI membrane translocation. These results indicate that membrane translocation of CAPRI and its function as a GAP protein are sequential and yet independent.

### Inhibitor for the GPCR-mediated adaptation regulates the sensitivity of a eukaryotic cell

Motile Escherichia coli (*E. coli*) provides the simplest and the best understood model of chemosensing system that is also mediated by chemoreceptors (59, 60). Modification of the chemoreceptors and interactions among the chemoreceptors have been proposed for a robust response, a precise adaptation, and a high sensitivity of chemoreceptors in bacteria (61–63). However, the molecular mechanism by which to control the GPCR-mediated adaption and the sensitivity of a eukaryotic cell in chemotaxis are not fully understood. A local excitation and global inhibition (LEGI) model explained how a eukaryotic cell responds to chemoattractant stimuli with a wide-concentration range, achieves adaptation, and establishes the intracellular polarization (10, 64). Different from *E. coli*, chemoattractant stimulation induces a persistent activation of heterotrimeric G protein in *Dictyostelium* (6, 11, 26), indicating that adaptation occurs downstream of GPCR/G protein in the eukaryotic cell. We constructed a detailed model to explain adaptation and chemosensing of *Dictyostelium* cells (15, 65), which led us to uncover the essential role of Ras inhibitors in the GPCR-mediated adaptation (11, 16). Recently, it has been shown that reducing the PI(3,4)P2 level on the plasma membrane increases Ras activity and enhances the excitability in *D. discoideum* (18, 19) and that active Ras plays a role in basal locomotion and in chemotaxing through very shallow gradients (24). We also detected a higher basal activity of Ras and Rap1 in *capri^kd^* cells. These reports inspired us to test the chemotaxis capability of both CTL and *capri^kd^* cells using wide concentration-range gradients of various chemoattractants (Fig. 6 and Fig. S10). In response to fMLP gradients, CTL cells chemotaxed better than *capri^kd^* cells did in gradients at high concentrations (> 100 nM), while they displayed similar chemotaxis capability to *capri^kd^* cells in mild gradients (10 nM of fMLP). In weak gradients (< 1 nM), *capri^kd^* cells chemotaxed better than CTL cells did. Similar chemotaxis behavior was also observed when CTL and *capri^kd^* cells were exposing to gradients of IL-8. Interestingly, *capri^kd^* cells sensed and responded to weak gradients of fMLP and IL-8 at the concentration that are subsensitive to CTL cells due to a higher sensitivity of *capri^kd^* cells (Fig. 7 and Fig. 8). Thus, our study provides evidence that human neutrophils locally recruits Ras inhibitor, CAPRI, to regulate Ras adaptation and their sensitivity. CAPRI enables human neutrophils to chemotax through a higher-concentration range of chemoattractant gradients by lowering its sensitivity and by mediating the GPCR-mediated adaptation.

### Multiple RasGAP proteins might be involved in Ras deactivation in chemoattractant-induced adaptation of neutrophils

In the present study, we have shown that CAPRI mediates chemoattractant-induced Ras adaptation and sensitivity of neutrophils. Human neutrophils without CAPRI fail in chemoattractant-induced adaptive Ras activation; have significantly increased phosphorylation of AKT, GSK3α/3β, and cofilin; demonstrate excessive actin polymerization; and exhibit a subsequent defect in chemotaxis in response to high-concentration gradients. However, in response to a low-concentration (either 0.1 nM or 10 nM fMLP) stimulation, *capri^kd^* cells displayed transient Ras/Rap1 activation and actin polymerization, although they displayed prolonged, non-adaptive responses upon stimulation with higher concentrations (Fig. 1, Fig. 7, and Fig. 8). HL60 cells also expressed RASAL1 and RASAL2 (Fig. S2)(37), which also deactivate Ras (34, 39). In mouse and human neutrophils, there are other RasGAP proteins expressed in addition to CAPRI (Fig. S2). p120 GAP is present in both human and mouse neutrophils but absent in HL60 cells (66) and its role has been implicated in cell migration (67, 68). While we revealed a major role of CAPRI in mediating the GPCR-mediated Ras adaptation in HL60 cells, it is also crucial to determine potentially roles of other RasGAP proteins in Ras adaptation and chemotaxis in primary mammalian neutrophils.

## Supporting information

Manuscript with figures

## Acknowledgements

We thank Drs. Arjan Kortholt, Xinzhuan Su, and Peter Crompton for their critical reading of the article. This work was supported by the NIH Intramural Fund from the National Institute of Allergy and Infectious Diseases, National Institutes of Health.

## Materials and Methods

Detailed descriptions of cells, cell lines, cell culture, and differentiation, reagents and antibodies, plasmids and transfection of cells, immunoblotting of fMLP-mediated signaling components, Ras and Rap1 activation assay, actin polymerization assay, immunoprecipitation assay, imaging and data processing, TAXIScan chemotaxis assay and data analysis, and SiR-actin staining are in Supplementary information.

