## Supplementary material for "Ras Inhibitor CAPRI Enables Neutrophils to Chemotax Through a Higher-Concentration Range of Gradients": Manuscript with figures

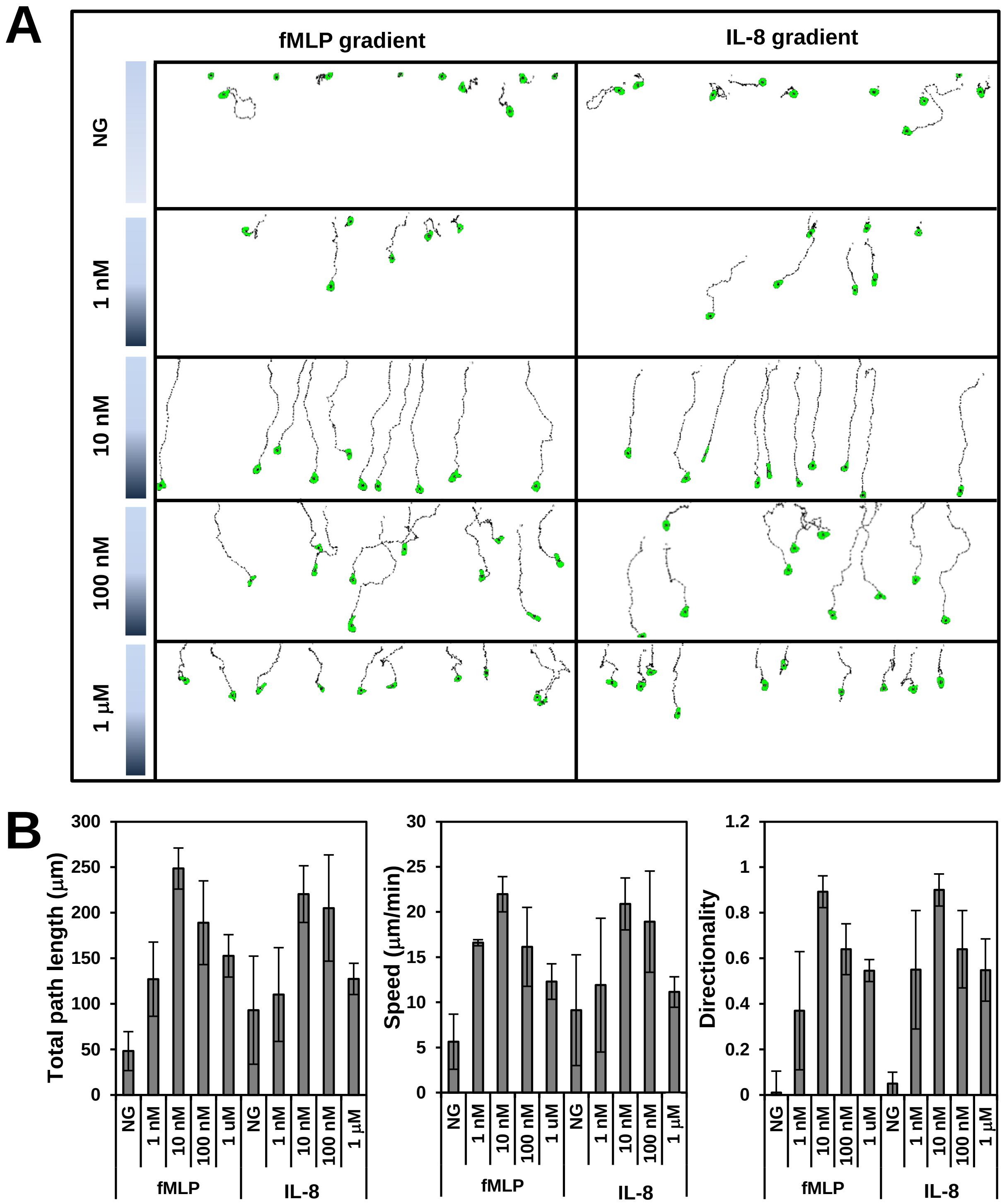
A
fMLP gradient
IL-8 gradient
NG
1 nM
10 nM
100 nM
1 µM
B
Total path length (µm)
Speed (µm/min)
Directionality
NG
1 nM
10 nM
100 nM
1 uM
1 µM
fMLP
IL-8

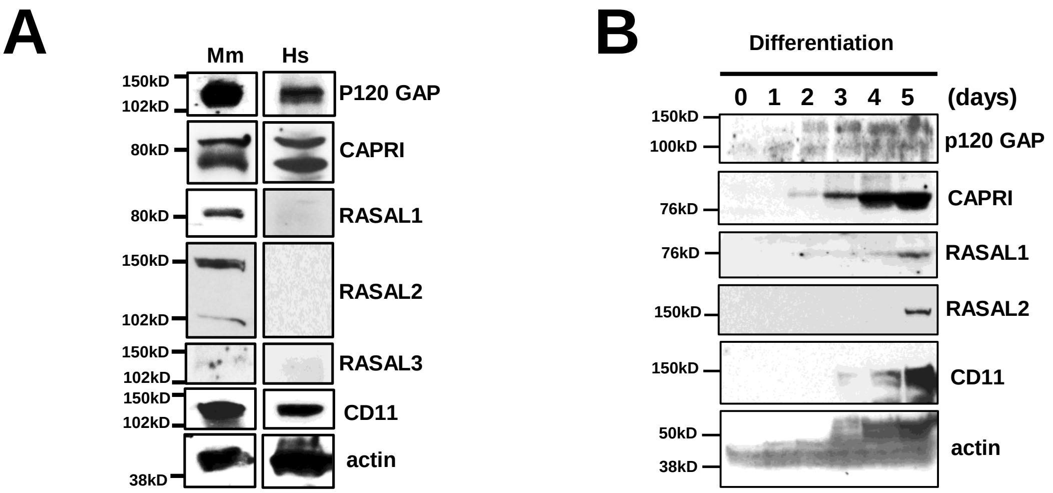
A
Mm
Hs
150kD
102kD
P120 GAP
80kD
CAPRI
80kD
RASAL1
150kD
102kD
RASAL2
150kD
102kD
RASAL3
150kD
102kD
CD11
38kD
actin
B
Differentiation
0 1 2 3 4 5 (days)
150kD
100kD
p120 GAP
76kD
CAPRI
76kD
RASAL1
150kD
RASAL2
150kD
CD11
50kD
38kD
actin

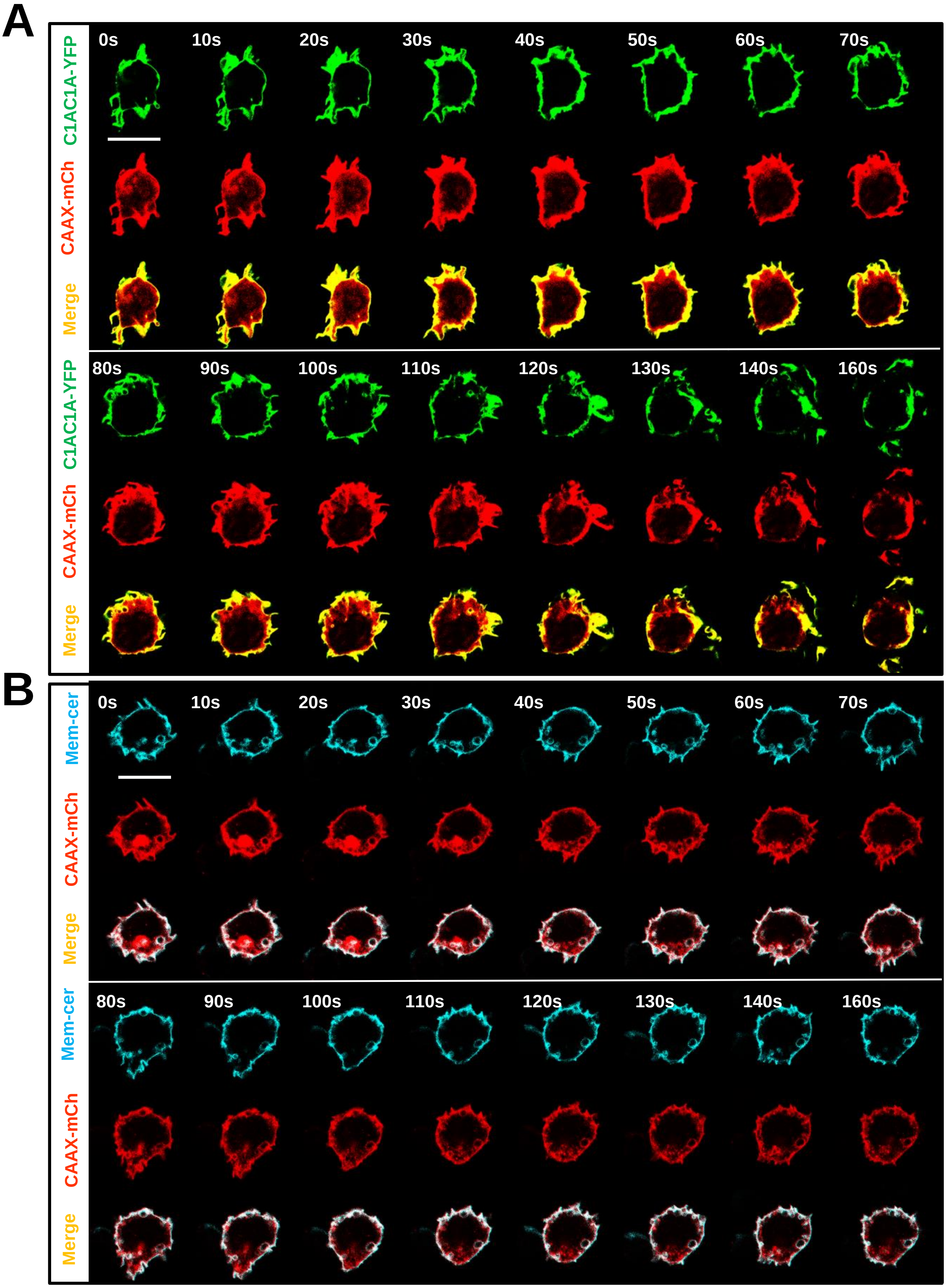
A
C1AC1A-YFP
CAAX-mCh
Merge
0s
10s
20s
30s
40s
50s
60s
70s
80s
90s
100s
110s
120s
130s
140s
160s
B
Mem-cer
CAAX-mCh
Merge

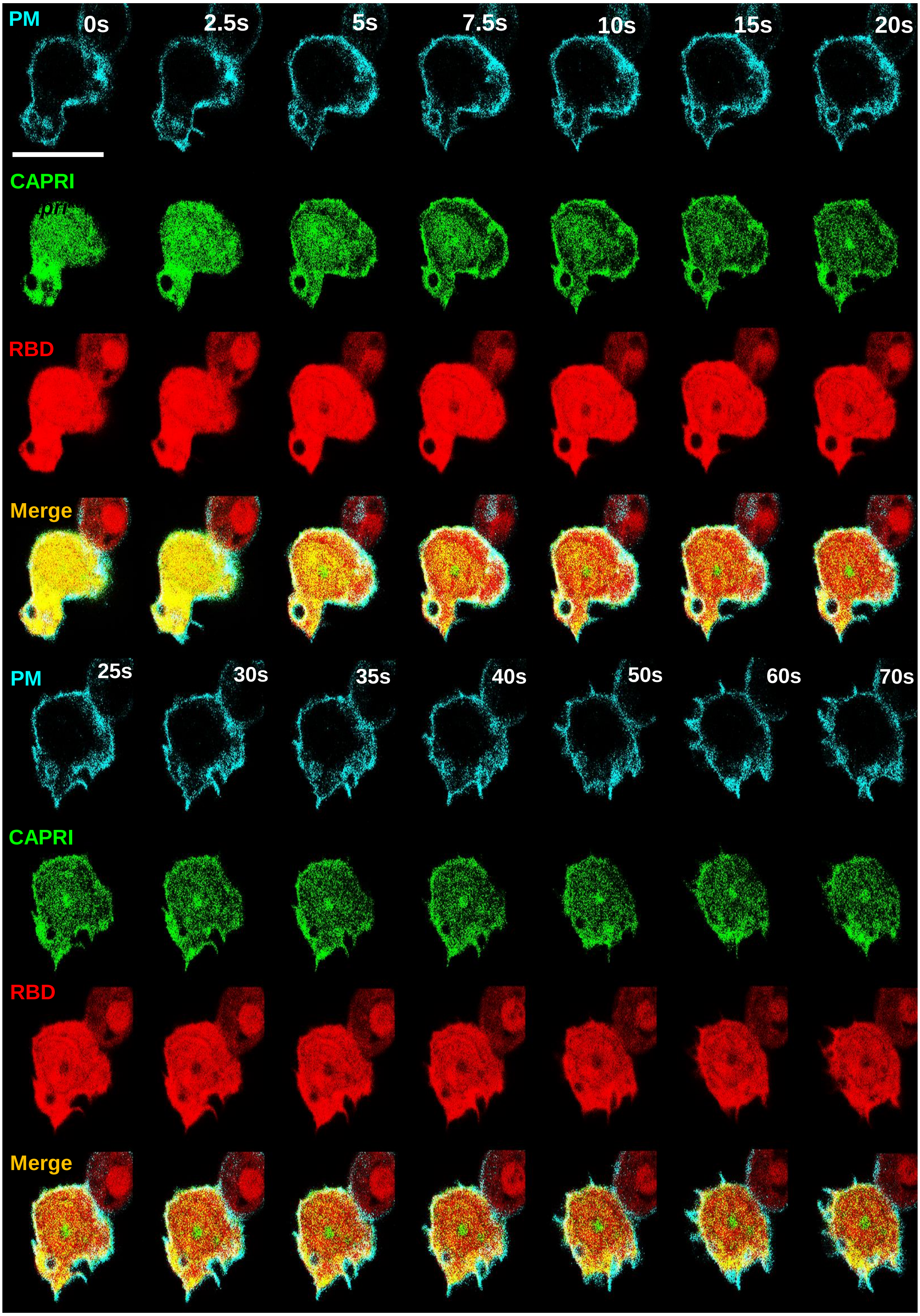
PM
0s
2.5s
5s
7.5s
10s
15s
20s
CAPRI
RBD
Merge
PM
25s
30s
35s
40s
50s
60s
70s
CAPRI
RBD
Merge

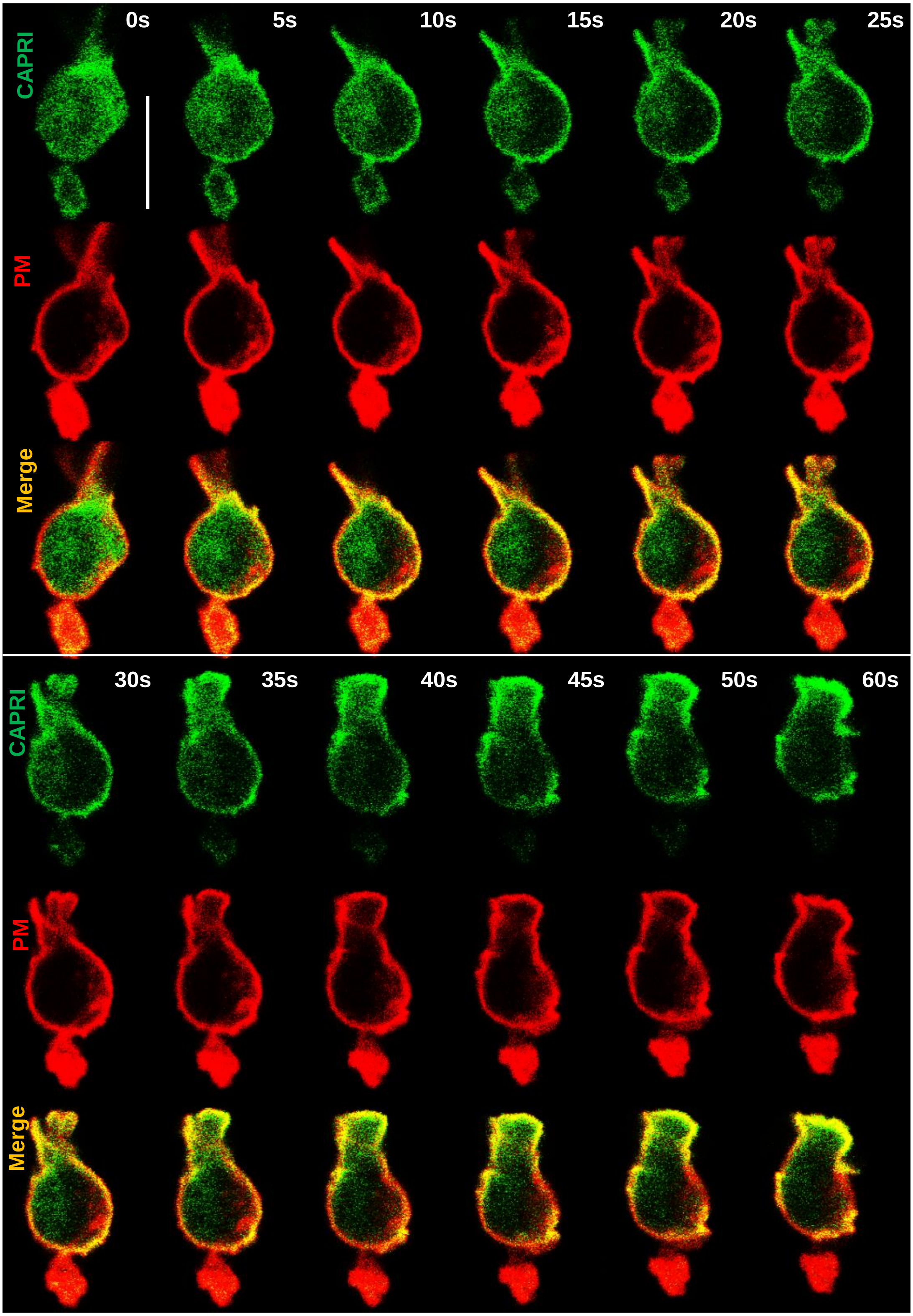
0s
5s
10s
15s
20s
25s
CAPRI
PM
Merge
30s
35s
40s
45s
50s
60s
CAPRI
PM
Merge

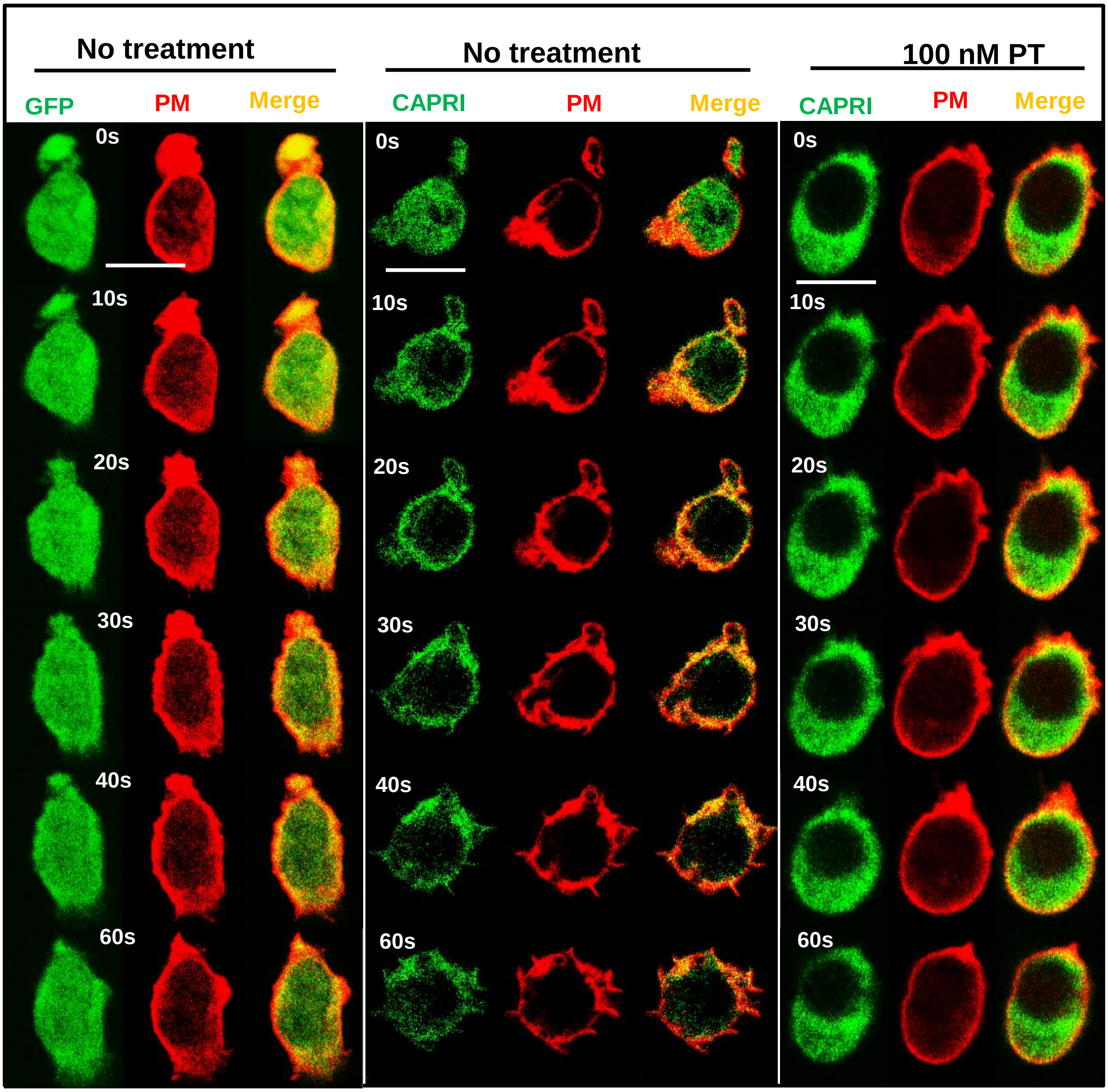
No treatment
GFP
PM
Merge
No treatment
CAPRI
PM
Merge
100 nM PT
CAPRI
PM
Merge
0s
10s
20s
30s
40s
60s
0s
10s
20s
30s
40s
60s
0s
10s
20s
30s
40s
60s

```
Hs NF1    LTRDLDQASMEAVVSLLAGLPLQPEEGDGVELM-EA------KSQLFLKYFTLFMNLLND    1116
Hs CAPRI  YPRWNET------------FEFELQEGAMEALCVEAWDWDLVSRNDFLGKV--VIDVQRL    222
Mm CAPRI  YPRWNET------------FDFELEKGASEALLVEAWDWDLVSRNDFLGKV--AVNVQRL    222
            *  :             : :: ::*    *  **      . : **  .   ::: .

Hs NF1    CSEVEDESAQTGGRKRGMSRRLASLRHCTVLAMSNLLNANVDSGLMHSIGLGYHKDLQTR    1176
Hs CAPRI  RVVQQE---------------------------EGWFRLQPDQSKSRRHDEGNLGSLQLE    255
Mm CAPRI  CSAQQE---------------------------EGWFRLQPDQSKSR-QGKGNLGSLQLE    254
              ::                           .. :. : *..  :  . *   .** .

Hs NF1    ATFMEVLTKILQQGTEFDTLAETVLAD-RFERLVEL---VTMMGDQGELPIAMALANVVP    1232
Hs CAPRI  ----------------VRLRDETVLPSSYYQPLVHLLCHEVKLGMQGPGQLIPLIEETTS    299
Mm CAPRI  ----------------VRLRDETVLPSVCYQPLVQLLCQEVKLGTQGPGRLIPVIEETTS    298
                          .    **** .  :: **.*    . :* **   :   : :..

Hs NF1    CSQWDELARVLVTLFDSRHLLYQLLWNMFSKEVELADSMQTLFRGNSLASKIMTFCFKVY    1292
Hs CAPRI  TECRQDVATNLLKLFLGQGLAKDFLDLLFQLELSRTSETNTLFRSNSLASKSMESFLKVA    359
Mm CAPRI  AECRQEVATTLLKLFLGQGLAKDFLDLLFQLELGRTSEANTLFRSNSLASKSMESFLKVA    358
           .  :::*  *:.** .: *  ::*  :*. *:  :.. :****.****** *   :**

Hs NF1    GATYLQKLLDPLLRIVITSSDWQHVSFEVDPTRLEPS--------------ESLEENQRN    1338
Hs CAPRI  GMQYLHGVLGPIINKVFEEKKYVE----LDPSKVEVKDVGCSGLHRPQTEAEVLEQSAQT    415
Mm CAPRI  GMRYLHGILGPIIDRVFEEKKYVE----LDPSKVEVKDVGCSGLHRPQTEAEVLEQSAQT    414
          *  **: :*.*::  *: ...: .    :**:::* .              * **:. :.

Hs NF1    LLQMTEKFFHAIISSSSEFPPQLRSVCHCLYQVVSQRFPQNSI-----GAVGSAMFLRFI    1393
Hs CAPRI  LRAHLGALLSALSRSVRACPAVVRATFRQLFRRVRERFPGAQHENVPFIAVTSFLCLRFF    475
Mm CAPRI  LRAHLVALLSAICRSVRTCPAIIRATFRQLFRRVRERFPNAQHQNVPFIAVTSFLCLRFF    474
          *      :: *:  *    *  :*:. : *:: * :***  .       ** * : ***:

Hs NF1    NPAIVSPYEAGILDKKPPPRIERGLKLMSKILQSIANHV---LFTKEEHMRPFNDFVKSN    1450
Hs CAPRI  SPAIMSPKLFHLRERHADARTSRTLLLLAKAVQNVGNMDTPASRAKEAWMEPLQPTVRQG    535
Mm CAPRI  SPAILSPKLFHLRERHADARTSRTLLLLAKAVQNIGNMDTPVSRAKESWMEPLQPTVRQG    534
          .***:**    : :::   * .* * *::* :*.:.*       :**  *.*::  *:..

Hs NF1    FDAARRFFLDIASDCPTSDAVNHSLSFISDGNVLALHRLLWNNQE--KIGQYLSSNRDHK    1508
Hs CAPRI  VAQLKDFITKLVDI--------------EEKDELDLQRTLSLQAPPVKEGPLF----IHR    577
Mm CAPRI  VAQLKDFIMKLVDI--------------EEKEELDLQRALNSQAPPVKEGPLF----IHR    576
              : *: .:..               .: : * *:* *  :    * *  :     *:
```

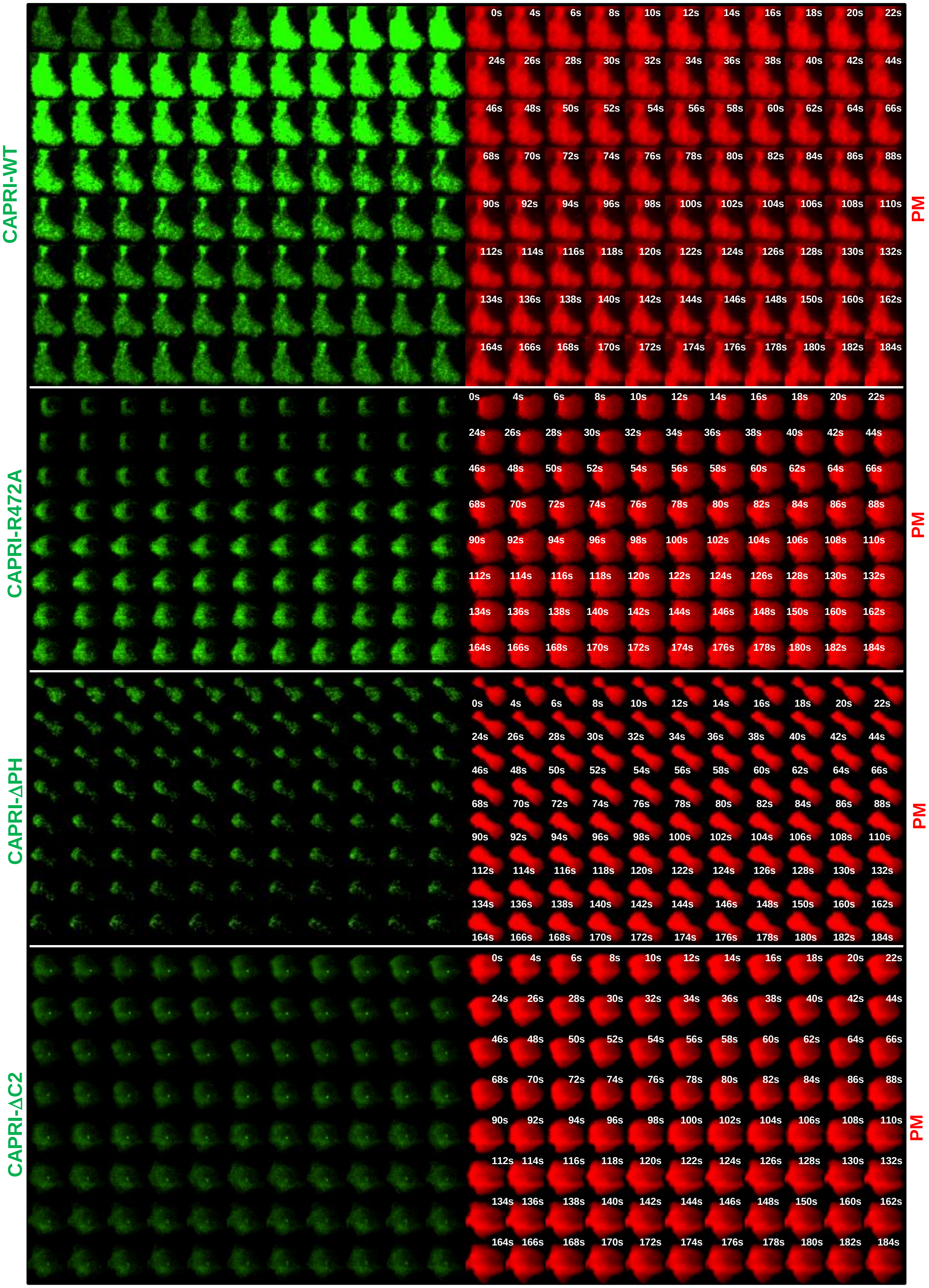
CAPRI-WT
0s 4s 6s 8s 10s 12s 14s 16s 18s 20s 22s
24s 26s 28s 30s 32s 34s 36s 38s 40s 42s 44s
46s 48s 50s 52s 54s 56s 58s 60s 62s 64s 66s
68s 70s 72s 74s 76s 78s 80s 82s 84s 86s 88s
90s 92s 94s 96s 98s 100s 102s 104s 106s 108s 110s
112s 114s 116s 118s 120s 122s 124s 126s 128s 130s 132s
134s 136s 138s 140s 142s 144s 146s 148s 150s 160s 162s
164s 166s 168s 170s 172s 174s 176s 178s 180s 182s 184s
PM
CAPRI-R472A
0s 4s 6s 8s 10s 12s 14s 16s 18s 20s 22s
24s 26s 28s 30s 32s 34s 36s 38s 40s 42s 44s
46s 48s 50s 52s 54s 56s 58s 60s 62s 64s 66s
68s 70s 72s 74s 76s 78s 80s 82s 84s 86s 88s
90s 92s 94s 96s 98s 100s 102s 104s 106s 108s 110s
112s 114s 116s 118s 120s 122s 124s 126s 128s 130s 132s
134s 136s 138s 140s 142s 144s 146s 148s 150s 160s 162s
164s 166s 168s 170s 172s 174s 176s 178s 180s 182s 184s
PM
CAPRI-ΔPH
0s 4s 6s 8s 10s 12s 14s 16s 18s 20s 22s
24s 26s 28s 30s 32s 34s 36s 38s 40s 42s 44s
46s 48s 50s 52s 54s 56s 58s 60s 62s 64s 66s
68s 70s 72s 74s 76s 78s 80s 82s 84s 86s 88s
90s 92s 94s 96s 98s 100s 102s 104s 106s 108s 110s
112s 114s 116s 118s 120s 122s 124s 126s 128s 130s 132s
134s 136s 138s 140s 142s 144s 146s 148s 150s 160s 162s
164s 166s 168s 170s 172s 174s 176s 178s 180s 182s 184s
PM
CAPRI-ΔC2
0s 4s 6s 8s 10s 12s 14s 16s 18s 20s 22s
24s 26s 28s 30s 32s 34s 36s 38s 40s 42s 44s
46s 48s 50s 52s 54s 56s 58s 60s 62s 64s 66s
68s 70s 72s 74s 76s 78s 80s 82s 84s 86s 88s
90s 92s 94s 96s 98s 100s 102s 104s 106s 108s 110s
112s 114s 116s 118s 120s 122s 124s 126s 128s 130s 132s
134s 136s 138s 140s 142s 144s 146s 148s 150s 160s 162s
164s 166s 168s 170s 172s 174s 176s 178s 180s 182s 184s
PM

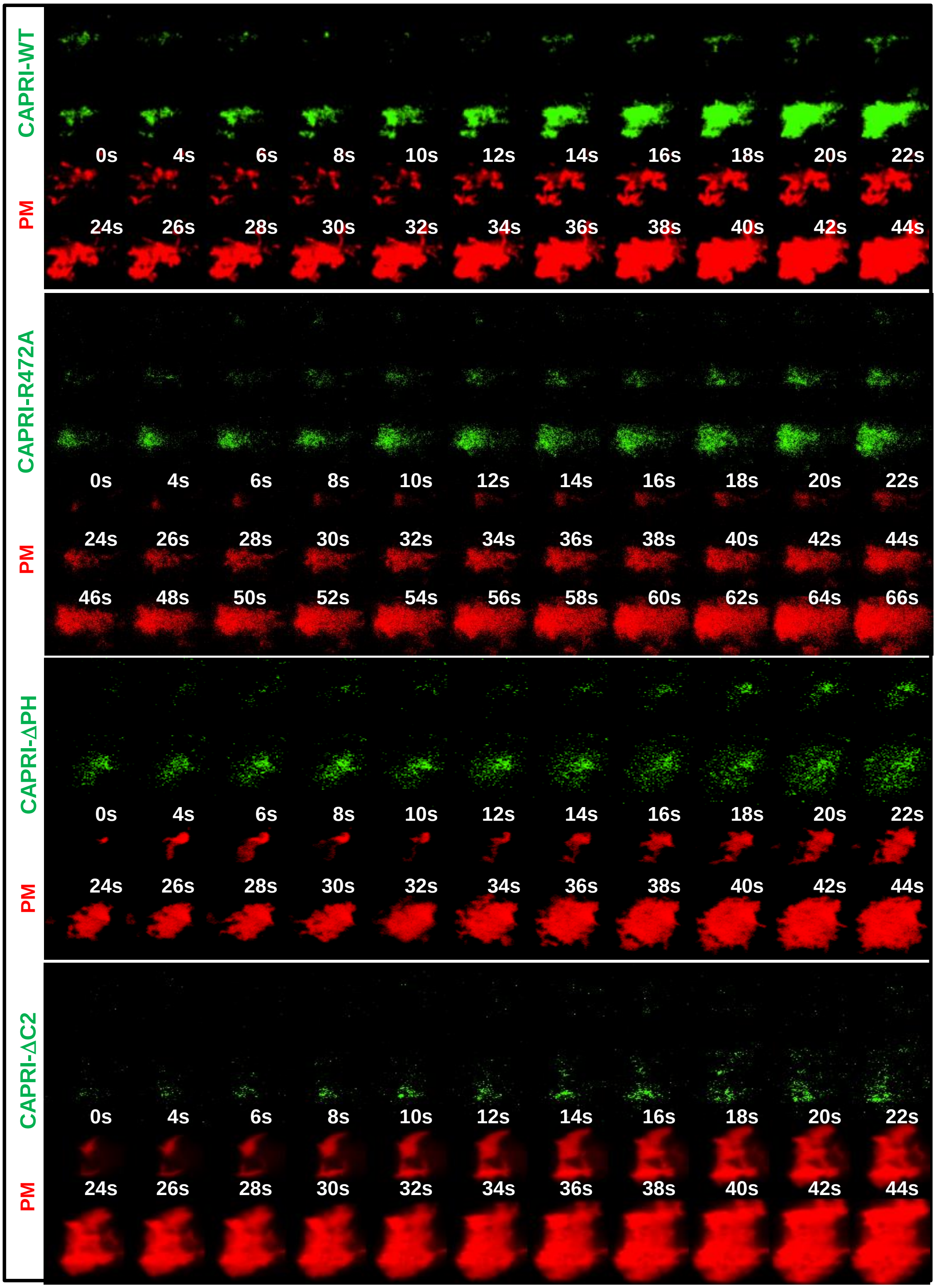
CAPRI-WT
PM
0s 4s 6s 8s 10s 12s 14s 16s 18s 20s 22s
24s 26s 28s 30s 32s 34s 36s 38s 40s 42s 44s
CAPRI-R472A
PM
0s 4s 6s 8s 10s 12s 14s 16s 18s 20s 22s
24s 26s 28s 30s 32s 34s 36s 38s 40s 42s 44s
46s 48s 50s 52s 54s 56s 58s 60s 62s 64s 66s
CAPRI-ΔPH
PM
0s 4s 6s 8s 10s 12s 14s 16s 18s 20s 22s
24s 26s 28s 30s 32s 34s 36s 38s 40s 42s 44s
CAPRI-ΔC2
PM
0s 4s 6s 8s 10s 12s 14s 16s 18s 20s 22s
24s 26s 28s 30s 32s 34s 36s 38s 40s 42s 44s

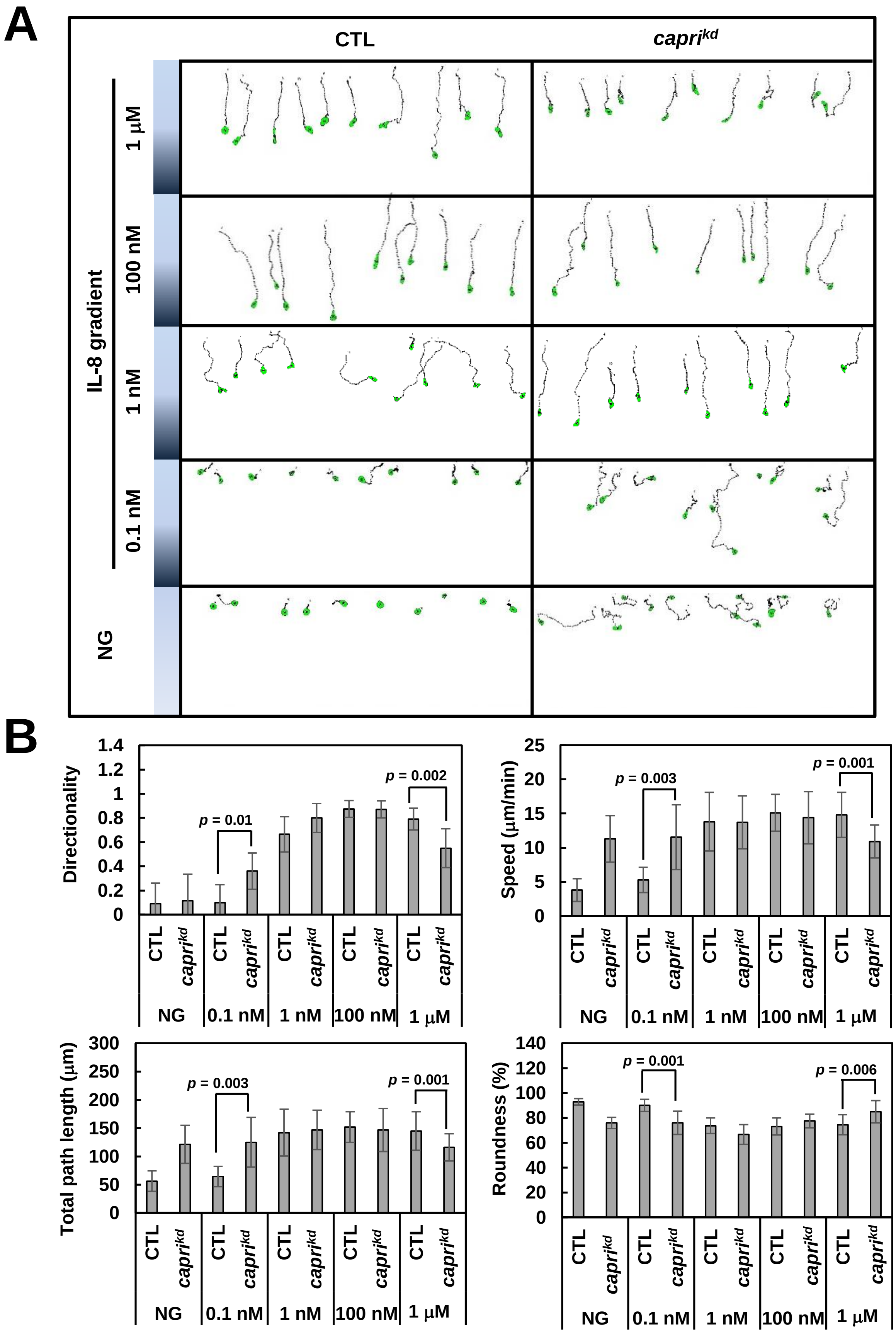
A
CTL
capri^kd
IL-8 gradient
1 μM
100 nM
1 nM
0.1 nM
NG
B
Directionality
p = 0.01
p = 0.002
Speed (μm/min)
p = 0.003
p = 0.001
Total path length (μm)
p = 0.003
p = 0.001
Roundness (%)
p = 0.001
p = 0.006
CTL
capri^kd
NG
0.1 nM
1 nM
100 nM
1 μM

**Supplemental Information**

**Supplemental Figure Legends:**

**Fig. S1.** GPCR-mediated chemotaxis of HL60 cells in response to fMLP and IL8 gradients with an enormous concentration range. (A) Montages show the travel path of chemotaxing HL60 cells in response to a wide range of fMLP and IL-8 gradients. (B) Chemotaxis behaviors measured from **A** and described as three parameters: directionality, where 0 represents random movement and 1 represents straight movement toward the gradient; speed, defined as the distance that a cell's centroid moves as a function of time; total path length, the total distance a cell has traveled.

**Fig. S2.** Mammalian neutrophils and HL60 cells highly express CAPRI. (A) The expression pattern of RasGAP proteins in human and mouse neutrophils. (B) The expression pattern of RasGAP proteins in human neutrophil-like (HL60) cells during DMSO-induced differentiation. Aliquots of cells, which were differentiated in the RPMI culture media with 1.3 % DMSO for the indicated days as described in the session of Materials and Methods, were mixed with SDS loading buffer, sonicated, and then subject to western blot detection of interested proteins of interests with their specific antibodies.

**Fig. S3.** Three plasma membrane markers colocalize in HL60 cells before and after chemoattractant fMLP stimulation.  (A) Colocalization of two membrane markers **(**C1AC1A-YFP and CAAX-mCherry) in HL60 cells. Cell expressing two plasma membrane (PM) markers, C1AC1A-YFP (green) and CAAX-mCherry (red), was stimulated with fMLP at time 0s. During the period of imaging, C1AC1A-YFP colocalizes with in the cells before and after fMLP stimulation. (B) Colocalization of two membrane markers **(**mem-cerulean and CAAX-mCherry) in HL60 cells. Cell expressing two plasma membrane (PM) markers, mem-cerulean (cyan) and CAAX-mCherry (red), was stimulated with fMLP at time 0 s. Mem-cerulean colocalizes with CAAX-mCherry before and after fMLP stimulation. Scale bar = 10 μm in **A** and **B**.

**Fig. S4.** *capri*$^{kd/OE}$ cell displayed little membrane translocation of active Ras probe RBD-RFP while displayed a strong membrane translocation of CAPRI. The *capri*$^{kd/OE}$ cell expressing PM marker (cyan), CAPRI-tGFP (green), and active Ras probe RBD-RFP (red) was stimulated with 1 μM fMLP at time 0 s and displayed a clear membrane translocation of CAPRI-tGFP and little membrane translocation of RBD-RFP. Scale bar = 10 μm.

**Fig. S5.** IL-8 induces membrane translocation of CAPRI in HL60 cells. HL60 cell expressing CAPRI-GFP (green) and PM marker (red) was stimulated with IL8 (100 ng/ml) at time 0 s. Scale bar = 10 μm.

**Fig. S6.** Heterotrimeric G protein inhibitor pertussis toxin blocks membrane translocation of CAPRI. Montage shows that the membrane translocation of CAPRI was inhibited in the HL60 cell treated with 100 nM pertussis toxin (PT). HL60 cells expressing PM marker (red) and GFP alone (left panel), or CAPRI-tGFP (CAPRI, green) without (middle panel) or with 100 nM PT (right panel) treatment were then stimulated with 1 μM fMLP at time 0 s.

**Fig. S7.** The identification of R-fingers in Hs NF1, Hs CAPRI, and Mm CAPRI. R-finger is a key feature of GAP domains. R473 on Hs CAPRI and R472 of Mm CAPRI are corresponding to R1391 of Hs NF1, a well-characterized arginine residue for the arginine finger. Amino acid sequences were obtained from https://www.ncbi.nlm.nih.gov/pubmed/ and aligned using software ClustalW2 provided from https://www.ebi.ac.uk.

**Fig. S8.** Membrane translocation of CAPRI wild-type (WT) or mutants in response to fMLP stimulation monitored using total internal reflection fluorescent (TIRF) microscopy. HL60 cells expressing tGFP-tagged WT or mutants of CAPRI (CAPRI-WT, -R472A, -ΔPH, or -ΔC2, green) and PM marker (red) were monitored by TIRF microscopy and stimulated with 1 μM fMLP at time 10 s.

**Fig. S9.** CAPRI is recruited to the adhesion sites of cell. Montage shows recruitment of CAPRI WT and its mutants to the adhesion sites during landing and spreading of HL60 cells. HL60 cells expressing CAPRI-tGFP (CAPRI, green) and PM marker (red) were monitored by TIRF microscopy.

**Fig. S10**. The concentration range of IL-8 gradients in which $capri^{kd}$ cells sense and chemotax is upshifted. (A) Montages show the traveling path of chemotaxing CTL or $capri^{kd}$ cells in response to a dilution series of IL8 gradients. Cells were examined with EZ-TAXIScan using an IL-8 dilution series. (B) Chemotaxis behavior measured as three parameters, including directionality, speed, and total path length, is shown. The definition of the three parameters is detailed in the figure legend of **Fig. S1B**. Student's *t* test was used for the *p* value.

**Supplemental Movie legends**

**Videos S1-S3.** fMLP-induced Ras activation in the control (CTL), $capri^{kd}$, and $capri^{kd/OE}$ HL60 cells. Cells expressing Ras-GTP binding domain (RBD) of human Raf1 tagged with RFP (RBD-RFP, red) and PM marker (green) in CTL (**Video S1**) and $capri^{kd}$ (**Video S2**) HL60 cells. $capri^{kd/OE}$ cells (**Video S3**) expressing PM marker (mem-cerulean, light blue), RBD-RFP (red), and tGFP-CAPRI (green). Cells were monitored using fluorescence microscopy and stimulated with fMLP (1 μM) at time 0 s.

**Videos S4-S7.** fMLP-induced membrane translocation of tGFP-tagged WT or mutants of CAPRI was monitored using fluorescence microscopy. Cells expressing tGFP-tagged WT, R472A, ΔC2, or ΔPH of CAPRI (green) and PM marker (red) (**Video S4-S7**, respectively) were stimulated with fMLP at 0 s.

**Videos S8-S9**. fMLP-induced $PIP_3$ production in CTL (**Video S8**) and *capri$^{kd}$* (**Video S9**) HL60 cells were monitored using fluorescence microscopy. $PIP_3$ production was monitored by the membrane translocation of PIP3 biosensor PH-GFP (PH domain of human AKT1 tagged with GFP, green). Cells expressing PH-GFP (green) and PM marker (red) were stimulated with 1 μM fMLP at time 0 s.

**Videos S10-S11**. fMLP-induced Rap1 activation in CTL (**Video S10**) and *capri$^{kd}$* (**Video S11**) HL60 cells were monitored using fluorescence microscopy. Rap1 activation was monitored by the membrane translocation of active Rap1 probe, GFP-$RBD_{RalGDS}$, (Rap1-GTP binding domain of Ral-GDS tagged with GFP, green). Cells expressing GFP-$RBD_{RalGDS}$ (green) and PM marker (red) were stimulated with 1 μM fMLP at time 0 s.

**Videos S12-S13**. fMLP-induced actin polymerization in CTL (**Video S12**) and *capri$^{kd}$* (**Video S13**) cells. Actin polymerization was monitored by the membrane translocation of F-actin probe, GFP-F-tractin. Cells expressing GFP-Ftractin (green) and PM marker (red) were stimulated with 1 μM fMLP at time 0 s.

**Videos S14-S15.** F-actin distribution in chemotaxing CTL (**Video S14**) and *capri$^{kd}$* (**Video S15**) cells. Cells were stained with the actin filament probe SiR-actin (red) 30 min prior to the experiments and exposed to a gradient originated from 1 μM fMLP source. To visualize the fMLP gradient, fMLP (1 μM) was mixed with fluorescent dye Alexa 488 (green) at a final concentration of 1 μg/ml.

**Videos S16-S19**. The responsiveness of CTL (**Videos S16** and **S18**) and *capri*$^{kd}$ (**Videos S17** and **S19**) cells in response to 0.1 nM or 10 nM fMLP stimulation, respectively. Cells expressing GFP-Ftractin (green) and PM marker (red) were stimulated with fMLP at the indicated concentration after the first scan of the movies. Cellular response was visualized by the membrane translocation of GFP-F-tractin.

**Supplementary information:**

**Purification of human and mouse neutrophils.** Mouse neutrophils were purified from bone marrow as previously published (1). Briefly, one end of the hind legs of C57/B6 mice was cut open and bone marrow was collected by centrifugation at 8600 × *g* for 10 s. Red blood cells were removed with ACK buffer (Lonza) at room temperature. Cells were counted and subjected to a neutrophil isolation procedure using a mouse neutrophil isolation kit (Miltenyi Biotec, order No. 130097658). Collected neutrophils were stored in PBS supplemented with 2% FBS and 2 mM EDTA at 4°C. The purity of neutrophils was determined by staining the cells with mouse Gr-1/Ly-6G PE-conjugated antibody and FITC-conjugated CD11b antibody (R&D systems). The purity was above 95%.

Human neutrophils were purified from human blood as previously reported (2). Whole blood (150 ml) was drawn from healthy volunteers at the Blood Bank of the National Institutes of Health. Coagulation was prevented by heparin. The majority of the red blood cells were removed by dextran (0.2 g/l; GE Healthcare, Pittsburgh, PA) sedimentation (30 min, room temperature). Upper phases containing white blood cells were collected and washed twice in phosphatebuffered saline (PBS). Cells were resuspended in PBS and layered on top of a five-step Percoll gradient (65, 70, 75, 80, and 85%; Sigma-Aldrich) in 15-ml conical tubes. After centrifugation (800 × *g*, 20 min, room temperature), the 70/75/80% Percoll layers containing granulocytes were collected and washed twice in PBS. The collected neutrophils were resuspended in PBS, the cell concentration was determined, and the cells were kept at room temperature until use. Cell viability was determined by trypan blue dye extrusion, and the results were >98% viable neutrophils. The purity of the preparations was determined by Wright-Giemsa staining, and the yield was >95% neutrophil granulocytes.

**Immunoprecipitation assay.** Briefly, cells expressing turbo-GFP (tGFP) alone or tGFP tagged WT or mutants of CAPRI were starved with RPMI 1640 medium containing 25 mM HEPES at 37℃ for 3 hours as previously described (2). Cells were then collected and resuspended at $2×10^7$ cells/ml and then stimulated with a final concentration of 10 μM fMLP. Aliquots of cells at indicated time points were lysed with 10 ml immunoprecipitation buffer (IB, 20 mM Tris, pH 8.0; 20 mM $MgCl_2$; 10 % glycerol; 2 mM $Na_3VO_4$; 0.25 % NP40; and complete 1X EDTA-free proteinase inhibitor) with or without 1 mM GTPγS for 30 min on ice. Cell extracts were centrifuged at 16,000 × *g* for 10 min at 4 ℃. Supernatant fractions were collected and incubated with IgG beads pre-conjugated with anti-tGFP at 4 ℃ for 2 hours. Beads were washed four times with immunoprecipitation buffer, and proteins were eluted by boiling the beads in 50 μl SLB. The supernatants and eluted samples were subjected to western blot detection of the indicated proteins.

**Ras and Rap1 activation assay**. The procedure was as previously reported (3-6). Briefly, HL60 cells were starved with RPMI 1640 medium containing 25 mM HEPES at 37 ℃ for 3 hours. Cells were then collected and resuspended at $2×10^7$ cells/ml and set on ice for 10 min. Then the cells were transferred to a medical cup, shaken at a speed of 200 rpm for 3 min, and then stimulated with fMLP at the indicated final concentrations. At the indicated time points after stimulation, 100 μl of the cells was taken and mixed with IB and 10 μl of the cells was mixed with 10 μl SLB. The mixtures with IB buffer were incubated on ice for 30 min and then centrifuged at 100,000 × *g* at 4 ℃ for 30 min. The supernatants were incubated with agarose beads conjugated with RBD (active Ras binding domain of human Raf1) (Cytoskeleton, Inc. Denver, CO) or $RBD_{RalGDS}$ (Rap1-GTP binding domain of human RalGDS) (abcam, Cambridge, UK) at 4 ℃ for 2 hours. The agarose beads were washed three times with IB. The protein on the beads was eluted by mixing with 25 μl SLB. The eluted proteins were subjected to western blot detection of the indicated proteins to determine the active Ras/Rap1 protein. Aliquots of cells mixed with SLB were subject to sonication to determine the total Ras protein.

**Actin polymerization assay.** The protocol of G and F actin measurement mostly followed the instructions of the "G-actin /F-actin in Vivo Assay Kit" from Cytoskeleton Inc. (Denver, CO) (3). Briefly, 100 μl aliquots of cells at the indicated time points before and after 10 μM fMLP stimulation were mixed with 500 μl LAS2 buffer containing 1X LAS1 buffer (provided in the kit), 10 mM ATP and 1X proteinase inhibitor cocktail (Roche Life Science, Indianapolis, IN). The mixture of samples was homogenized using a 200 μl pipet tip and incubated at 37˚C for 10 min. Next, the samples were centrifuged at 350 × g for 10 min to pellet unbroken cells. A total of 400 μl supernatant of each sample was transferred to a centrifugation tube and centrifuged at 100,000 × *g* at 4˚C for 1 hour. This step is to pellet the F-actin and leave the G-actin in the supernatant. Supernatants were transferred to prelabeled 1.8 ml centrifugation tubes and mixed with 400 μl SDS loading buffer. To the pellet, 400 μl actin depolymerization buffer was added and incubated on ice for 1 hour to allow actin depolymerization to occur. Pipette up and down several times every 15 minutes to help pellet resuspension. 400 μl SDS loading buffer was added to each pellet sample. 10 μl of each sample was subjected to SDS-PAGE and Western blot to detect and analyze G- and F-actin amounts at given time points for the cell lines.
